# mamp-ml: A deep learning approach to epitope immunogenicity in plants

**DOI:** 10.1101/2025.07.11.664399

**Authors:** Danielle M. Stevens, David Yang, Tatiana J. Liang, Tianrun Li, Brandon Vega, Gitta L. Coaker, Ksenia Krasileva

## Abstract

Eukaryotes detect biomolecules through surface-localized receptors, key signaling components. A subset of receptors survey for pathogens, induce immunity, and restrict pathogen growth. Comparative genomics of both hosts and pathogens has unveiled vast sequence variation in receptors and potential ligands, creating an experimental bottleneck. We have developed mamp-ml, a machine learning framework for predicting plant receptor-ligand interactions. We leveraged existing functional data from over two decades of foundational research, together with the large protein language model ESM-2, to build a pipeline and model that predicts immunogenic outcomes using a combination of receptor-ligand features. Our model achieves 73% prediction accuracy on a held-out test set, even when an experimental structure is lacking. Our approach enables high-throughput screening of LRR receptor-ligand combinations and provides a computational framework for engineering plant immune systems.

## Main Text

Surface-localized pattern recognition receptors (PRRs) bind and respond to different biomolecules to initiate signal transduction, constituting a major class of innate immune receptors in plants and animals. In plants, PRRs are classified as receptor kinases (RKs) or receptor proteins (RPs) based on the presence or absence, respectively, of an intracellular kinase domain. These receptors can be further subdivided by phylogenetic placement and ectodomain structure (1–2). The leucine-rich repeat (LRR) ectodomain is the most abundant in both plants and animals with hundreds of receptors encoded in a single species, yet most remain uncharacterized with unknown ligands (1, 3). At a population level, the number of paralogs and alleles reveals further sequential variation, with growing evidence of the impact it can have on pathogen recognition and colonization (4–7).

PRR-binding ligands include microbe-associated molecular patterns (MAMPs), such as peptides from bacterial flagellin or polysaccharides derived from bacterial or fungal cell walls, damage-associated molecular patterns (DAMPs) derived from the plant cell, various endogenous plant-derived peptides and hormones as well as pathogen effector proteins (8). Pathogen and plant-derived peptides constitute a major class of PRR ligands and, upon receptor binding, require a conserved co-receptor(s) for signaling activation. Interactions between receptor and protein ligand were classically described as agonistic (immune-inducing) or antagonistic (immune-blocking) (9–10). However, characterization of diverse epitope variants has revealed alternate outcomes. Some non-immunogenic epitopes prevent receptor binding and promote pathogen colonization (8). Other epitopes act as weakly immunogenic deviant peptides, which display bimodal immune outcomes (5, 7, 11). Functional receptor identification and transfer across species can transform plant resilience and therapeutic development, but these studies rely upon large functional screens, both costly in terms of time, personnel, and resources, limiting the potential to transform discovery into implementation (4, 11–14). We postulate that the ability to computationally predict immunogenic outcomes of ligands with their associated receptor in a high-throughput manner would transform plant engineering and sustainable agriculture.

Recent advances in machine learning have enabled the effective use of large-scale protein sequence databases to learn biologically meaningful information. Protein language models (pLMs), such as Evolutionary Scale Modeling 2 (ESM-2), are trained using masked sequence modeling, where certain amino acids are hidden and the model learns to predict them based on the surrounding sequence. This approach enables the model to capture statistical patterns and evolutionary relationships within protein sequences (15–16). Trained on millions of protein sequences, these models capture rich representations and long-range patterns, capturing information on both secondary and tertiary structure and function, and can be readily transferred to downstream tasks with limited and/or unbalanced labeled data (17–19). Here, we present the ESM-2 pLM fine-tuned to predict immunogenic outcomes of LRR-type PRRs (LRR-PRRs) and protein ligands on just their sequence alone, providing the ability to *in silico* screen a large sequence space and prioritize novel functional variants efficiently. We foresee our work will advance the discovery, characterization, and engineering of receptor-protein ligand interactions.

### Digitizing two decades of functional receptor-protein epitope research

To curate the training and validation datasets, we manually mined existing literature for characterization of LRR receptor-protein ligand interactions. Currently, 11 plant LRR-PRR receptors are known to interact with small protein ligands (also termed epitopes) between 10 and 55 amino acids in length from diverse organisms (Fig. 1A). We focused on well-established epitopes that lack internal post-translational modifications and their associated plant receptors. While this primarily included defense-associated receptors in the phylogenetic XII subfamily, we additionally included receptors of the XI subfamily that recognize host-derived damage-associated molecular patterns and phytocytokines, plant epitopes that amplify immune responses (20–21). These peptides are derivatives of proteins thought to be cleaved and released via plant proteases (22–23). For both receptor classes, signaling outcomes can be classically described as immunogenic, non-immunogenic, or weakly immunogenic, which displays intermediate outputs and partial signaling induction, based on an array of standard immune assays (Fig. 1A).

**Fig. 1.**
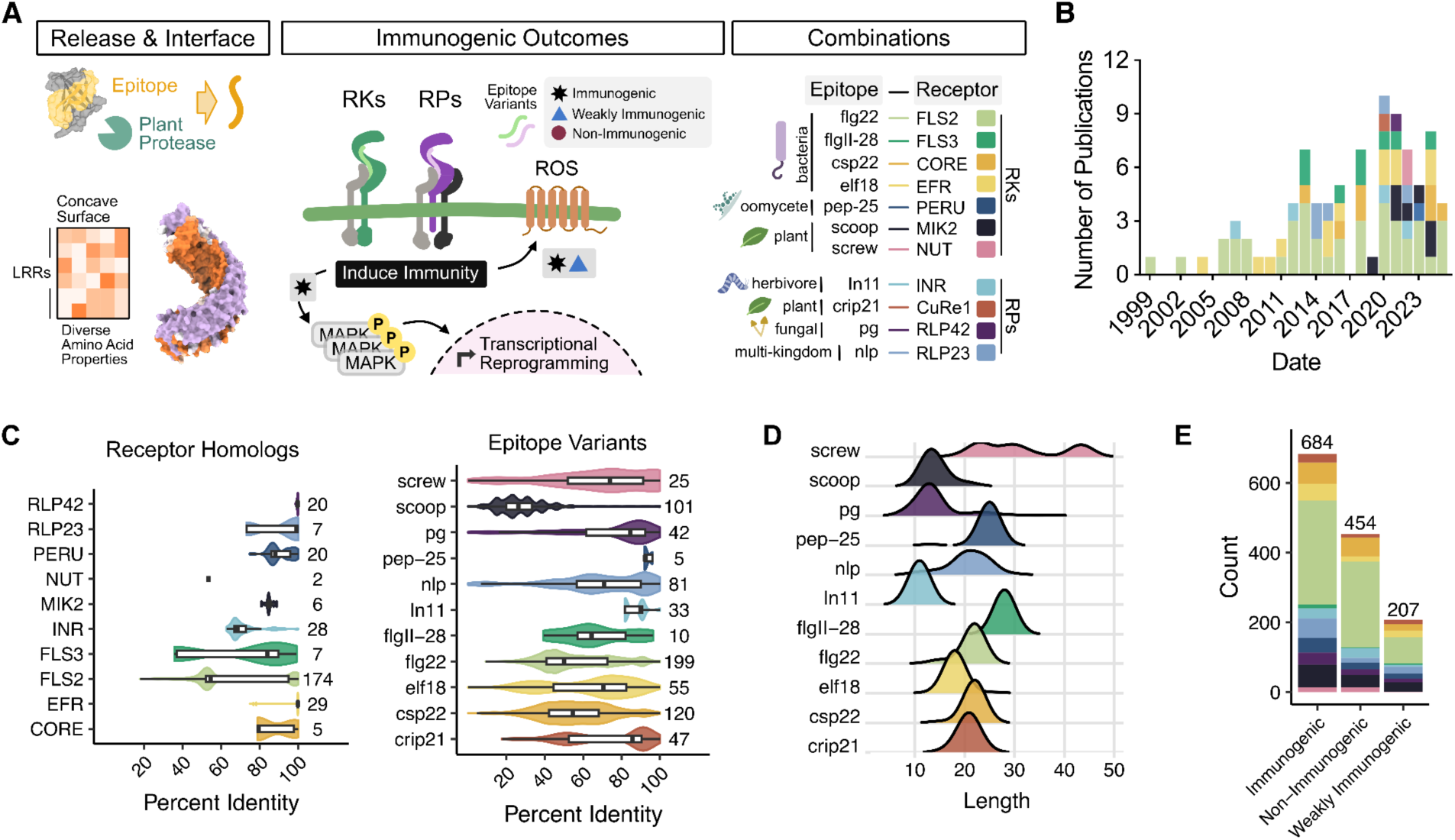
Digitizing two decades of foundational functional receptor-epitope ligand research. **(A)** Many epitopes are released from proteins through plant proteases. The leucine-rich repeat (LRR) concave surface represents a binding surface that designates the ligand profile spaces. Upon primary receptor binding, co-receptor(s) create a complex that initiates a signaling cascade and associated immune response. Diverse epitopes have shown alternated outcomes, including weak immunogenicity that leads to bimodal immune outputs. We focus on well-characterized receptor-epitope combinations such as flg22-FLS2 and flgII-28-*FLAGELLIN SENSING RECEPTOR 3* (FLS3), cold shock protein csp22-CORE, and elongation factor Tu elf18-*ELONGATION FACTOR RECEPTOR* (EFR), herbivore-associated inceptin in11-*INCEPTIN RECEPTOR* (INR), fungal polygalacturonases (pg)-*RECEPTOR-LIKE PROTEIN 42* (RLP42), parasitic plant glycine rich protein crip21-CuRe1, oomycete transglutaminase pep-25-*PEP13 RECEPTOR UNIT* (PERU), and multi-kingdom ligands necrosis and ethylene-inducing peptide 1 (Nep1)-like proteins (nlp)-*RECEPTOR LIKE PROTEIN 32* (RLP32) and phytokines scoops-*MALE DISCOVER 1-INTERACTING RECEPTOR-LIKE KINASE* (MIK2) and screw-*PLANT SCREW UNRESPONSIVE RECEPTOR* (NUT). **(B)** The number of publications with data available across 11 receptor-epitope pairs. **(C)** Receptor (left) and epitope (right) sequence diversity based on all-by-all percent identity comparisons. The total number of sequences assessed are on the right of each plot. **(D)** Distribution of sequence length across all epitopes of interest. **(E)** Distribution of receptor-epitope interaction data (n = 1339) collected for model training and validation based on their immunogenic outcomes.

Across 74 published studies spanning from 1999 to 2025, we obtained 1,339 unique interactions from 11 receptors and 91 plant species (Fig. 1B) (Supplemental Table 1). We collected additional training data of a well-characterized PRR-ligand pair, *FLAGELLIN SENSING RECEPTOR 2* (FLS2) receptor–bacterial flagellin-derived flg22 peptide, based on previous analyses (Fig. S1) (7, 11). For reference, CTNIP-HSL3 is the same as SCREW-NUT. By mining naturally occurring and synthetic receptor-epitope sequences, we find considerable sequence diversity, in particular for ligands, based on all-by-all global protein similarity (Fig. 1C). Ligands additionally display notable length variation within and across different epitopes (Fig. 1D). These receptor-ligand combinations were categorized by their immunogenic outcomes based on the assay, controls, and previous description leading to 684 immunogenic, 454 non-immunogenic, and 207 weakly-immunogenic pairs (Fig. 1E). The diversity of data collected, as well as previous comparative genomics of plant receptors and protein ligands, highlights the large sequential space possible and the need for *in silico* functional screening (7, 13).

AlphaFold3 has previously been suggested as a computational screening method for prediction receptor-ligand interactions, based on using the protein-protein interaction metric, ipTM (interface predicted TM-score) (11, 24–25). By comparing these metrics in past studies with another receptor-epitope interaction collected here, we found FLS2-flg22 and PEPR-pep, both which have solved structures, performed favorably when using the standard ipTM cutoff of 0.8 with AlphaFold3 (hereafter AF3) (11). Other receptors such as known variants of LRR-RK *ELONGATION FACTOR RECEPTOR* (EFR)-elf18 and LRR-RK *CORE*-csp22 displayed low model confidence metrics, where known immunogenic and non-immunogenic interactions could not be distinguished (3) (Fig. S2A-B). We hypothesize that AF3 incorrectly attempts to create secondary structure in the unstructured ligand and misaligns the side-chains, leading to poor ipTM scores (Fig. S2C). Therefore, due to limited defense-associated LRR-PRRs structures that bind to small protein ligands, a computational approach that does not rely on structural context is required (3). Since there is considerable functional data on PRRs and their cognate ligands, we used this data to train the foundational pLM ESM-2 model for immunogenic outcomes.

### Integrating LRR information with pLM architecture

Our goal was to build a generalizable model that could be used across receptors with different LRR repeat numbers. While it is well-established that the concave side of the LRR ectodomain serves as the ligand binding site (26–27), accurate prediction of LRR repeats has been challenging due to variance in both repeat length and composition. Therefore, we built a pipeline that extracts the LRR ectodomain from full-length receptor sequences using AlphaFold2 (AF2) and a computational dimensionality reduction-based LRR annotator method, LRR-annotation (28). AlphaFold2 enabled adequate globular protein prediction with average pLDDT scores above 80, indicative of confident structural prediction (Fig. 2A). This observation reflects the conserved solenoid structure of LRRs found across diverse domains of life (29–30). The structure-model aware dimensionality reduction method, LRR-annotation, transforms the 3D structure into a 2D space where the LRR forms a circular coil, to track the number of LRR repeats and precisely extract the ligand-binding LRR ectodomain (28). We find this method enables accurate extraction of LRR ectodomains across both RKs and RPs from diverse plant species (Fig. 2B). For some receptors, however, the LRR repeat number was underestimated yet still captured the ligand binding region (Fig. 2B, Fig. S3B). Overall, LRR-annotation correctly determines the number of LRR repeats for most receptors based on previous literature (Fig. 2B).

**Fig. 2.**
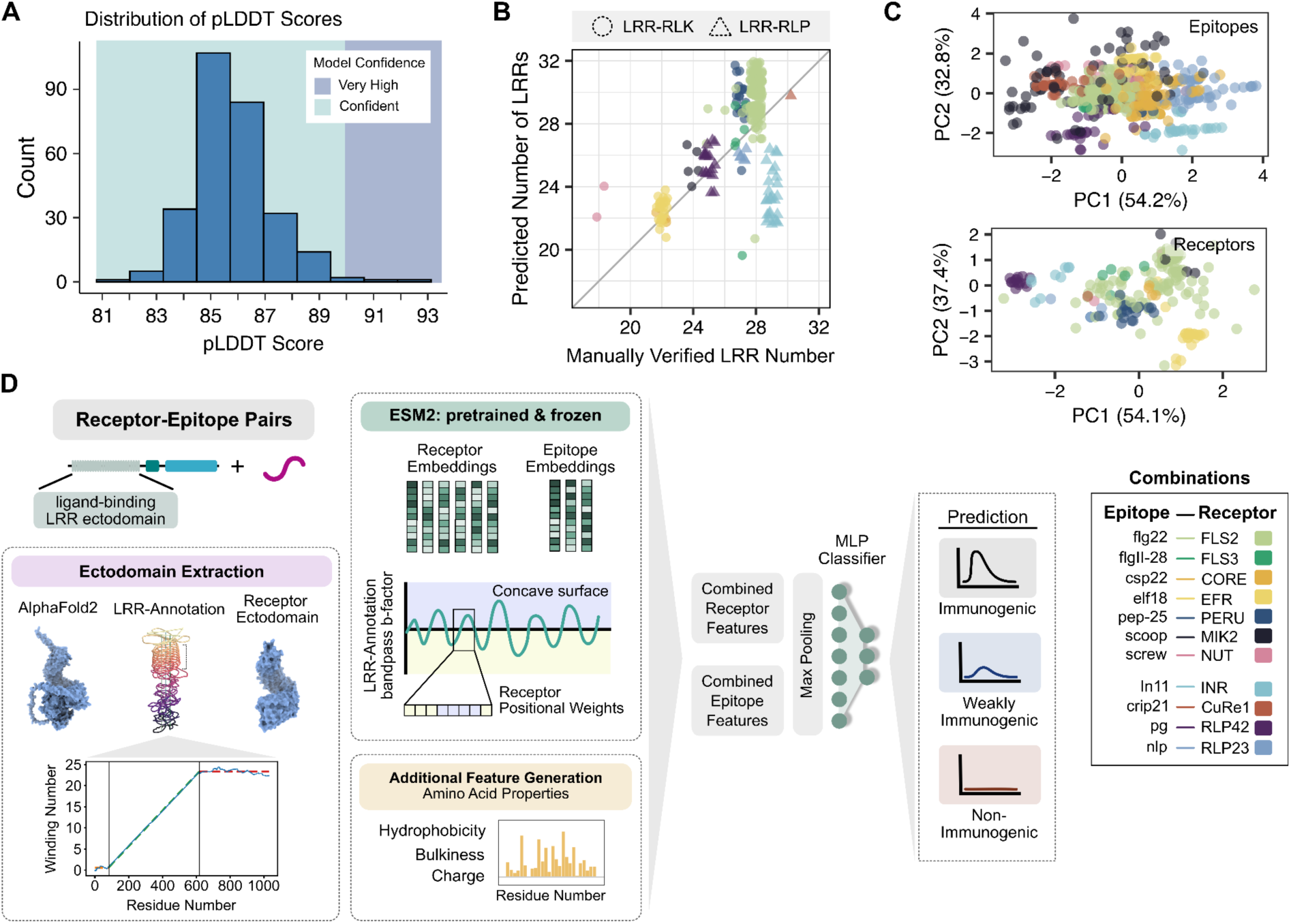
Integrating LRR-epitope information with pLM ESM-2 to generate mamp-ml, a fine-tuned model for predicting immunogenicity. **(A)** AlphaFold2 structural model metrics average pLDDT across all 290 receptor sequences. **(B)** Predicted number of LRRs by LRR-Annotation versus manually verified. **(C)** Principal component analysis of epitope (top) and receptor (bottom) chemical features including average amino acid bulkiness, charge, and hydrophobicity. **(D)** Model pipeline includes receptor preprocessing for precise LRR ectodomain extraction through AlphaFold2 and LRR-Annotation, and feature generation of amino acid properties for receptors and protein ligands. The paired final-layer receptor-epitope embeddings are extracted from pLM ESM-2, and the receptor weights are adjusted so residues along the inner concave surface are emphasized during model training. The embeddings in combination with amino acid properties are fine-tuned to classify for immunogenic, weakly immunogenic, and non-immunogenic outcomes.

Previous studies characterizing plant receptor-protein epitope interactions have shown that amino acid features such as bulkiness, charge, and hydrophobicity impact ligand recognition profiles (11, 13, 31–32). We recognized these could be used as distinct features to aid differentiating receptors and ligands (Fig. S4, Fig. S5). Using an unsupervised approach via a principal component analysis (PCA), we similarly find these features can be used to distinguish receptors and protein ligands (Fig. 2). To improve prediction accuracy between different receptor-ligand pairs, we sought to integrate these features into our predictive model.

Our approach, mamp-ml, uses the final layer of the 8-million-parameter ESM-2 model to embed both receptor and epitope protein sequences and integrate them with amino acid–level features for classifying immunogenicity (Fig. 2D). After applying layer normalization on the chemical features, we concatenated both sequence embeddings and amino acid property vectors for both receptor ectodomain and protein epitope. This combined matrix was then inputted into a multi-layer perceptron classifier. Since the inner concave surface of the receptor is essential for ligand binding, we used the bandpass b-factor values from LRR-annotation to adjust the weights for the receptor, with positive values (inner surface) receiving higher weights than negative values (outer surface) (Fig. S3) (11). To reduce overfitting, we applied dropout and L2 regularization. These techniques enabled the model to learn PRR receptor-ligand interactions and predict how amino acid mutations affect immunogenic outcomes. Our approach enables rapid screening of hundreds to thousands of receptor–ligand variants within minutes.

### Assessing a generalizable model for LRR-protein ligand interactions

To predict immunogenicity without structural information, we fine-tuned the protein language model ESM-2 using several architecture variants: a sequence-only version with embeddings and classifier head (SeqOnly), and a version incorporating amino acid chemical features, including bulkiness, charge, and hydrophobicity (ChemAll) (Fig. S7A). Our final model, mamp-ml, integrates these chemical features with positional weights from LRR-annotation. All three models were trained and evaluated using an 80-20% stratified split (1,071 training, 268 test combinations) based on immunogenicity, which indirectly stratified sequence diversity since immunogenic and non-immunogenic receptor-ligand pairs exhibit distinct sequence patterns and evolutionary relationships that correlate with their functional outcomes (Fig. 3A, Fig. S7).

**Fig. 3.**
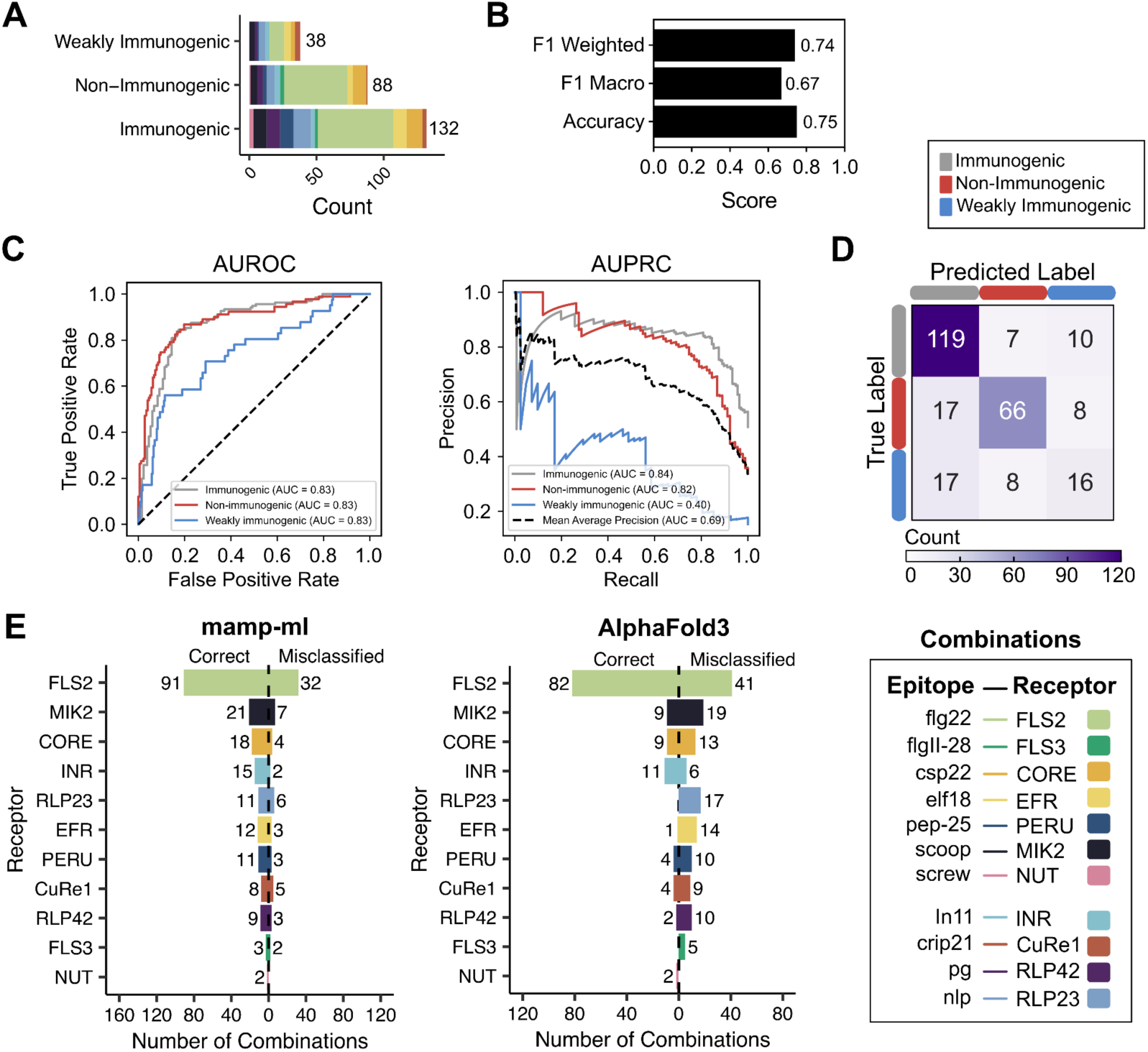
Performance of mamp-ml with validation data in comparison to AlphaFold3. **(A)** Validation data distributed by immunogenic outcomes. **(B-C)** Performance metrics of mamp-ml by (B) accuracy, F1 macro, F1 weighted, AUROC (Area Under the Receiver Operating Characteristic curve), and (C) AUPRC (Area Under the Precision-Recall Curve) for each labeled class. **(D)** Confusion matrix of validation data displays false positive and false negative predictions across all labeled classes. **(E)** Number of correct and misclassified predictions per receptor using mamp-ml (left) and AlphaFold3 with 0.8 ipTM as a cutoff (right).

Mamp-ml outperformed baseline approaches across all evaluation metrics, including F1-macro, F1-weighted, and accuracy (Fig. 3B, Fig. S7B). AUPRC analysis revealed that mamp-ml effectively distinguished immunogenic (0.84) and non-immunogenic (0.82) receptor-ligand combinations but struggled with weakly immunogenic cases (0.40) (Fig. 3C). Mamp-ml performs favorably on other metrics such as AUROC (0.83) and F1-weighted (0.74) (Fig. 3B-C). To confirm that the model learned genuine receptor-ligand interactions rather than spurious patterns, prediction scores dropped significantly when receptors and epitopes are shuffled in the training data (Fig. S7B-C). For SeqOnly, we did not observe the same drop in model accuracy, but we hypothesize that, due to the hypervariable nature of the LRR repeat, there was insufficient context to learn how to differentiate LRRs between different receptors (Fig. S7D). Given that ESM-2 models range from 8 million (8M) to 15 billion (15B) parameters, we additionally assessed the effect of pre-trained model size on prediction accuracy. Across four model sizes ranging from 8M to 650M parameters, we found that the smallest model (8M) provided accuracy without overfitting (Fig. S8).

Notably, mamp-ml had a higher misclassification rate of weakly immunogenic incorrectly predicted as immunogenic (Fig. 3D). We hypothesize this may be due to limited training data, which prevents the model from capturing subtle class differences or the quantitative nature of qualitative labels, leading to some immunogenic cases being mislabeled as weakly immunogenic (Fig. 2C, Fig. 3D). For three of the receptor-epitope pairs, we assessed if there was a particular signature that led to misclassification. There was no notable difference in global receptor or epitope sequence similarity (Fig. S9A, C). We observed a minor increase in misclassification in epitopes of slightly shorter length, providing a clear area for future model improvement (Fig. S9B). Despite this, mamp-ml performed favorably in predicting immunogenic outcomes across most receptor-epitope combinations assessed (Fig. 3E). We then ran our same validation dataset through AF3 to compare prediction potential. When an ipTM cutoff of ≥0.8 was used as a proxy for binding, AF3 could well predict the interactions of some receptor-epitope pairs such as FLS2-flg22 (Fig. 3E, Fig. S10). However, AF3 failed to do so for almost every other receptor-epitope combination evaluated, and those that were correctly predicted were likely an artifact of the model’s inability to differentiate these interactions, as most correct predictions were from non-immunogenic outcomes which display low ipTM scores (Fig. 3E, Fig. S10). Our results highlight that pLMs augmented with amino acid side chain information achieve greater prediction accuracy than computational structural prediction alone.

To assess the performance of mamp-ml on a new receptor-epitope combination, we evaluated our trained model on *PEP Receptor* (PEPR1/PEPR2), which recognizes the danger-associated molecular pattern known as peps (33). Using past research, we found a similar distribution of immunogenic, weakly immunogenic, and non-immunogenic outcomes as our training data (Fig. S11A). Both PEPR homologs and pep epitopes were sequentially diverse (Fig. S11B). We zero-shot predicted all 47 PEPR-pep combinations with both mamp-ml and AF3 to assess their ability to predict immunogenicity. We found mamp-ml performed poorly, only predicting 21% (10/47) combinations correctly, whereas AF3 could predict 62% (29/47) correctly (Fig. S11C-D). To improve prediction accuracy, mamp-ml requires training on some examples of the receptor-ligand pair to learn the associated LRR repeat pattern and its ligand profile. However, even with limited training data, as is the case with FLS3-flgII-28, notable prediction power is possible (Fig. 3E). As the discovery of receptors and ligands increases beyond those showcased here, the development of new features and models will enable improved prediction of new receptor-ligand interactions and receptor bioengineering.

### Mamp-ml predicts receptor-ligand interactions without structural context

To independently test mamp-ml on new functional datasets, we screened Solanaceous plants, including cultivated and wild relatives, for immunogenicity induced by the CORE receptor against a panel of 65 csp22 epitope variants (7, 34–35). The csp22 receptor CORE is an excellent candidate for independent validation as it is structurally conserved, spans multiple plant genera, and is sequentially diverse both within and outside of its epitope-binding LRR domain (Fig. 4A, Fig. S12). Likewise, csp22 is frequently a multicopy ligand that can be found across diverse pathogenic organisms with high sequence variation (Fig. 4B) (7, 13). We selected two species from the Solanaceous family that encoded a diverse CORE sequence to screen for immunogenic outcomes via Reactive Oxygen Species (ROS) production assay against the panel of 65 csp22 epitope variants (Fig. 4C). Using a similar range of classification based on the ROS magnitude, we categorized their interactions and used mamp-ml for immunogenicity prediction (Fig. 4D, Fig. S13).

**Fig. 4.**
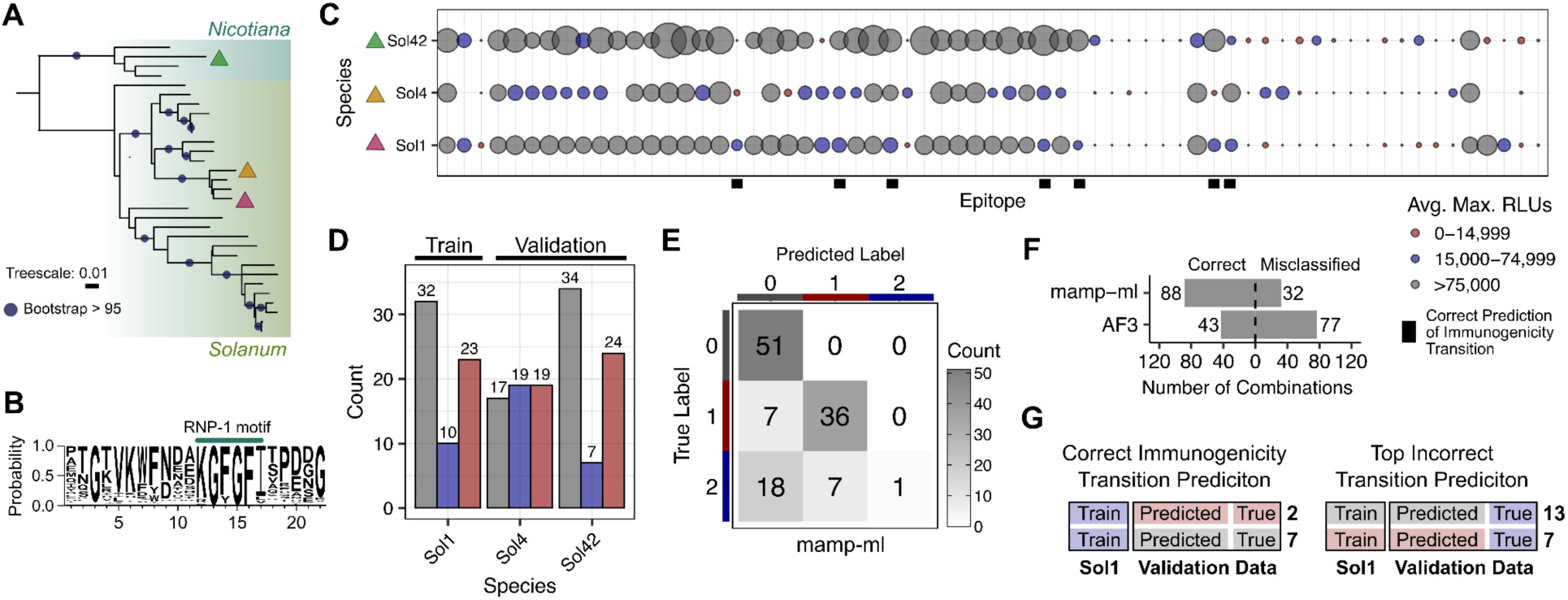
Using CORE-csp22 as independent validation, mamp-ml performs favorably with diverse receptor and ligand variants. **(A)** Maximum likelihood phylogenetic tree of 16 CORE homologs from the Solanaceous family. CORE receptors of interest are highlighted with triangles, where magenta is *S. lycopersicum* cv. Rio Grande (data included in training), green is *N. benthamiana* (validation data), and orange is *S. habrochaites* LA1353 (validation data). **(B)** Weblogo of 65 csp22 epitope variants used to survey immunogenic outcomes on Solanaceous species. **(C)** ROS screen for immunogenicity across csp22 epitope variants across three species which encode a CORE homolog. 200 nM peptide was tested with at least three plants per replicate and repeated two times. **(D)** Categorization of immune outcomes in (C). **(E)** Prediction of immunogenicity using mamp-ml displayed as a confusion matrix. Labels are the following: 0 = immunogenic, 1 = non-immunogenic, and 2 = weakly immunogenic. **(F)** Prediction of immunogenic outcomes by mamp-ml vs AF3 using an ipTM score of 0.8 as a cutoff. **(G)** Number of receptor-ligand pairs in our validation data where correct immunogenicity shifts were predicted and top outcomes where an incorrect shift was predicted.

Mamp-ml was primarily trained on CORE sequences from cultivated tomato, whereas the CORE homologs used to evaluate mamp-ml were from wild tomato *Solanum habrochaites* and the relative of tobacco, *Nicotiana benthamiana.* We found that mamp-ml predicted 73% (88/120) interactions correctly, whereas AF3 predicted only 36% (43/120) interactions correctly (Fig. 4E-F). Similar to our test set, all 43 correct predictions were due to low ipTM scores, which were found across all 120 CORE-csp22 combinations (Fig. S14). We then assessed the model’s ability to predict different immunogenicity states and found that mamp-ml was able to correctly predict either a transition to non-immunogenic or immunogenic interactions in our validation data (Fig. 4G). However, similar to our test data, mamp-ml struggled to predict weakly immunogenic outcomes in our validation data, especially when the training data was labeled as immunogenic or non-immunogenic (Fig. 4G).

In a recent study, a sequence-diverged, convergently evolved receptor from Citrus was discovered to recognize the csp22 ligand (13). Since CORE and SCORE bind the same ligand class, we postulated that mamp-ml may be able to predict immunogenic outcomes for SCORE-csp22 variants despite not being in the original training dataset. We mined Ngou et al. (13) for csp22 and SCORE variants, including orthologs, chimeras (LRR swaps), and single point mutations (AA substitutions), and categorized their immunogenic outcomes based on ROS induction (Fig. S15-17). Zero-shot prediction of each group displayed variable performance, with diverse orthologs displaying promising 42% (812/1957) prediction accuracy, a 21% improvement compared to our previous zero-shot receptor-ligand pair, PEPR-pep (Fig. 5A-B, Fig. S11, Fig. S18). Using SCORE-csp22 as a test case, we performed k-shot, n-classes training to assess how much data is required for a new receptor-ligand pair to improve prediction accuracy. We found notable improvement in AUROC, AUPRC, and F1 weighted scores with a k-shot size of 32 and above (Fig. 5C). When compared to our zero-shot predictions, we find between 22 and 56% improvement in prediction accuracy for 32 k-shot size (Fig. 5D). Even with larger k-shot sizes, prediction accuracy improvements are not remarkably different (Fig. 5D). We find mamp-ml can improve prediction after training with a minimal number of examples, highlighting its potential to expand to new receptor-epitope pairs.

**Fig. 5.**
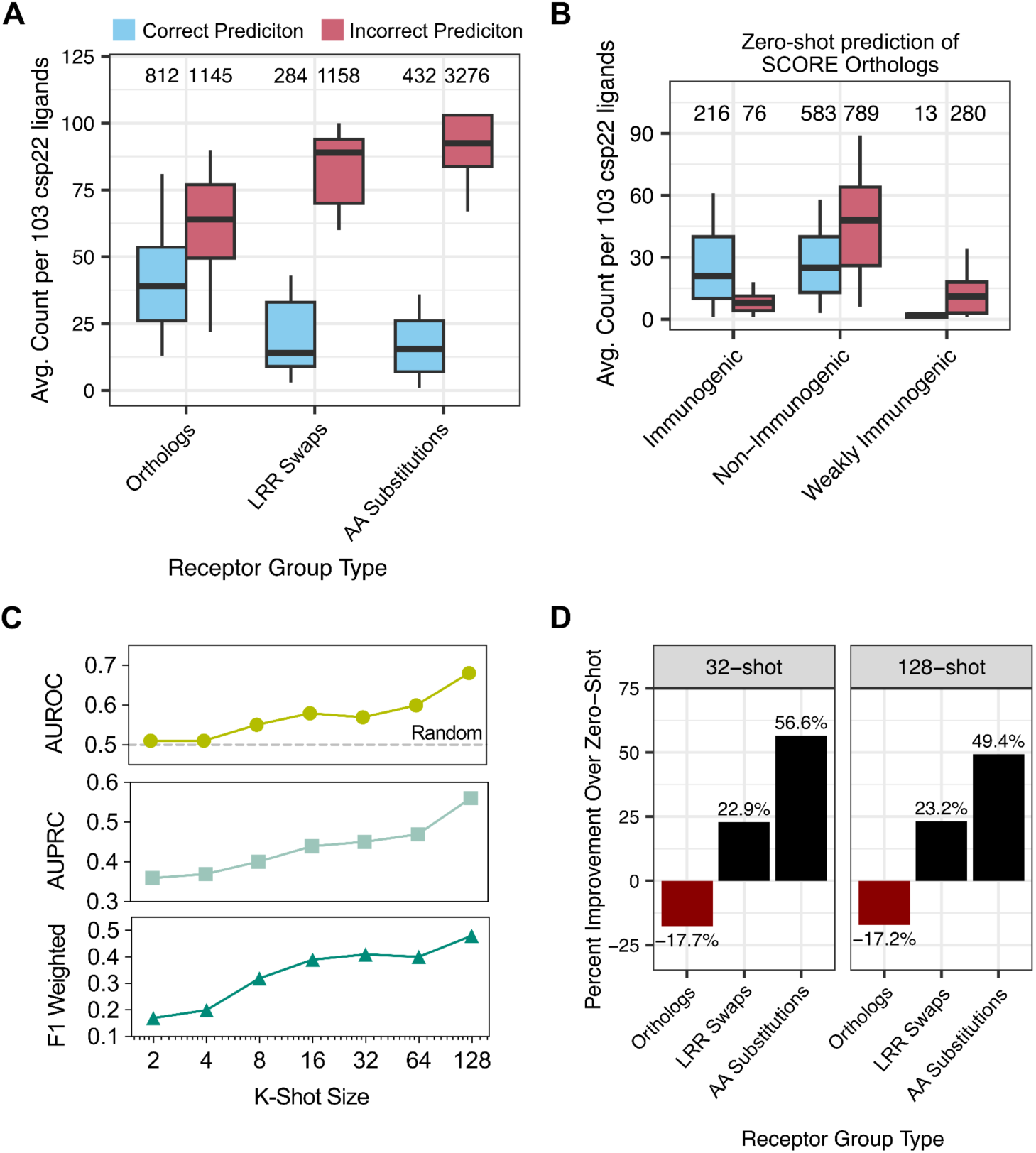
K-shot learning enables learning of new, convergently evolved receptor-epitope pair. **(A)** Zero-shot prediction of csp22-receptor SCORE and ligand variants, including orthologs, chimeras (LRR swaps), and single point mutations (AA substitutions), from Ngou et al. 2025 (13). For each receptor variant, 103 csp22 variant ligands were assessed. The total number of receptor-ligand combinations were plotted on top. **(B)** Average number of zero-shot correct predictions for SCORE orthologs in each immunogenic class. **(C)** Performance metrics of F1 weighted, AUROC, and AUPRC on SCORE-csp22 test data (n = 969) after training with different k-shot sizes. Hyperparameters for each k-shot size can be found in Supplemental Table 8. **(D)** Model prediction accuracy difference of 32- and 128-shot compared to zero-shot prediction across each receptor group type.

To make mamp-ml accessible, we implemented it on Google Colab and assessed runtime on different GPUs. Using the smallest GPU T4 (12.7 GB), which is also free, our csp22-CORE predictions took 55 minutes whereas on the largest GPU A100 (40 GB RAM), available on Colab Pro, runtime was cut down to 22 minutes (Supplemental Table 2). We then used mamp-ml on Colab to predict immunogenic outcomes on data from Ngou et al. 2025 (13) using the A100 GPU. To predict immunogenic outcomes on 2,163 combinations, which include 21 different receptor orthologs, took about one hour. For positive immunogenic prediction, a critical priority of new functional variant discovery, we achieved a zero-shot prediction accuracy of 74% (216/292) across SCORE orthologs (Fig. 5B). By splitting the receptor-ligands of interest over multiple GPUs, massive *in silico* screening is now possible narrowing the search space that needs to be tested in the laboratory for immunogenic interactions ten-fold. Our approach enables the prediction of immune outcomes despite lacking an experimentally validated structure, a common phenomenon for most defense-associated PRRs (3).

## Discussion

Eukaryotes deploy PRRs to sense and respond to diverse cues including pathogen attack, growth, and other physiological traits (21). Due to an evolutionary competition between hosts and colonizing organisms, both receptors and their associated ligands have expansive sequence diversity and gene copy number variations (2, 7). Until the emergence of affordable sequencing, most of immune receptor-epitope research focused on the biochemistry of single host-single epitope interactions. While this has provided a foundational understanding of host-microbe interactions, whole genome sequencing of both hosts and microbes has revealed considerable diversity and complexity, enabling new discoveries in signaling, pathogen evasion, and bioengineering (5–7, 11). For most interactions, functional outcomes and evolution of receptors and epitopes remain elusive. We have presented mamp-ml, the first computational framework for predicting interactions between LRR-PRRs and their protein ligands, enabling rapid *in silico* screening of receptor–epitope variants, and offering a scalable foundation for future immune receptor discovery and engineering.

Discovery of new ligands and their associated receptors has lagged considerably, despite the extensive diversity that exists (2, 13). Receptors were discovered using a range of experimental techniques including introgression lines, comparative genomics, and genome-wide association studies (GWAS), and most ligands are found using l biochemical approaches (4, 9, 13, 36). We show that protein language models can predict immunogenic outcomes of known receptors and significantly accelerate prioritization of new receptor variants, which is key for engineering broad recognition specificities. While mamp-ml outperformed the AF3 ipTM method with fair generalizability to ligands variants from diverse organisms and receptor homologs, it performed variably to new receptor-ligand pairs not seen during training (Fig. 5, Fig. S11), suggesting training data is key for improving prediction potential. Recently, Ngou et al. (13) reported the largest functional dataset available so far, enabling the discovery of SCORE, and allowing us to assess the testing power of mamp-ml. Promisingly, we discovered successful prediction of immunogenic outcomes in convergently evolved receptors such as SCORE, which share the same ligand as CORE (13) (Fig. 5A-B). It may be possible to use mamp-ml to discover additional convergently evolved csp22-receptors, which have previously been hypothesized in other species (37).

Mamp-ml marks a significant step beyond structure-based methods like AF3, with strong performance across diverse ligand variants and receptor homologs. LRR-PRRs represent a large protein family with an internal code or language within the concave surface. Despite considerable research since the discovery of the first receptor in the 1990s, our understanding of this code is insufficient. Our work highlights the potential expansion of machine learning methods and prediction approaches as new data become available, laying the groundwork for accelerated receptor discovery and engineering. Future work could explore additional fine-tuning on general receptor–ligand binding data, including structurally related innate immune receptors such as Toll-like receptors (TLRs) in animals, thereby generating computational solutions and therapeutic innovations for improved health across organisms.

## Acknowledgements

We thank Brian Su for helpful discussion regarding fine-tuning and architecture of large language models. AI was used for code debugging and documentation. This research was supported by the NIH Director’s New Innovator Award (1DP2AT011967-01) awarded to KK and NIH MIRA (2R35GM136402) awarded to GC.

## Author Contributions

DMS, KK, and GC conceived and designed the study. DMS mined for training data, performed bioinformatic analysis, built other subsequent model versions, helped collect CORE-csp22 data, and wrote the manuscript. DY built the SeqOnly model. TJL and VG collected CORE-csp22 validation data. TL collected additional training data. All authors were involved in writing and editing this manuscript.

## Data Accessibility

All data collected and AlphaFold models can be found in Zenodo (DOI: 10.5281/zenodo.15864665). The code base to build and evaluate mamp-ml can be found in GitHub (DanielleMStevens/mamp-prediction-ml). The production model and its implementation can be found in GitHub (DanielleMStevens/mamp-ml) and Colab.

## Methods

### Dataset curation for model training and validation

Receptor-ligand combinations were manually pulled from 74 manuscripts and categorized based on their immunogenicity, depending on the immune assay assessed, the relevant controls, and the authors’ previous description (Supplemental Table 1). Before model training could begin, we implemented a data processing pipeline that uses both AlphaFold2 (through a local ColabFold) (38) and the dimensionality reduction LRR annotator method (28) to model the receptor sequence and precisely extract the ectodomain containing just the LRR repeat domain. Specifically, each full-length receptor sequence was structurally modeled using colabfold_batch with –-num-models 1, and a custom Python script was used to parse LRR annotation output to extract the LRR domain sequence. Detailed code for computational analyses can be found in the following GitHub repository (DanielleMStevens/mamp-prediction-ml). To control for unequally distributed data, we performed an 80:20% split for training and validation data, stratifying either completely randomly between receptor and epitope sequences and using a random seed or stratified by immunogenic outcomes using scikit-learn (v1.6.1). To further test mamp-ml, we curated completely unseen receptor-epitope combinations from PEPR-pep literature and our own independent test data using new variants of receptor-epitope combination CORE-csp22 (Supplemental Table 3, 4).

### Model training

We utilize ESM-2, a transformer-based state-of-the-art protein language model (pLM) that ranges from 8M to 15B parameters (15). For most models and data splitting schemas, we started with the 30-layer, 150M parameter model (esm2_t30_150M_UR50D) trained on the UniRef50 database. All models were implemented in PyTorch (v2.6.0) and implementations of ESM-2 from the transformers package (v4.48.3). We used the pre-trained model as a feature extractor for each protein sequence and additionally fine-tuned the model weights for improved prediction accuracy. We additionally tested the impact of unfreezing 1, 2, and 3 layers of the esm2_t6_8M_UR50D model, and the hyperparameters selected were based on validation data set performance. The final model and associated weights were saved in GitHub: DanielleMStevens/mamp-ml.

For each receptor and epitope, the tokenizer function was used to return equal-length sequence embeddings. To infer long-range effects between receptor and epitope interactions and adjust for sequence length variation, the receptor and epitope were combined with a <eos> token, and additional padding was added to the length of the longest sequence of the batch. For the SeqOnly model, the embeddings were passed through a three-layer multilayer perceptron (MLP) for classification. For ChemAll, additional features were calculated including amino acid bulkiness, charge, and hydrophobicity using conversion values from the R package peptides (v4.4.1) (39). Before classification, the chemical features are processed through a FiLM (Feature-wise Linear Modulation) layer to transform each chemical feature vector into a sequential network, which is appended to the embeddings, which go through a max pooling step. These features are then similarly passed through an MLP for classification, reducing the dimension space from 1280 to 640 to a final projection of three classes. We used cross-entropy with logits loss to compute error for both SeqOnly and ChemAll, with the latter using both an L2 regularization lambda value of 0.01 and 0.2 dropout to limit overfitting. The final model, mamp-ml, is similar to ChemAll but additionally incorporates adjusting the positional weights in the feedforward based on the b-factor values calculated from the output of dimensionality reduction LRR annotation (28), where positive values the inner concave surface that binds with the peptide ligand have a weight of 2 and negative values which do not interact with the ligand have a weight of 0.5.

We tracked model performance using multiple standard metrics for classification, including F1 macro and weighted score, accuracy, AUROC, and AUPRC for each class. Due to the variable distribution of data, we additionally calculated an average AUPRC to capture performance across all classes. The best model was selected based on a combination of metrics. Models were trained for between 15 and 20 epochs and a batch size of 8 or 12. Model weights were optimized via backpropagation using the AdamW optimizer (40). Details of best best-performing hyperparameters for mamp-ml can be found in Supplemental Table 5.

### Computational analyses of receptors and protein ligands

To visualize how chemical features impact receptor and ligand uniqueness, a PCA was built similarly as described in Li et al. 2024 (11). Briefly, using the peptides R package (v4.4.1) (39), the average bulkiness, charge, and hydrophobicity using the Malavalin scale were calculated for both receptor ectodomain sequences and protein ligands (39). Each PCA was built after scale normalization using the prcomp function from the stats R package (v3.6.2).

To build weblogos of receptor and ligand sequences, unique variants were exported to a fasta file, and sequences were aligned using mafft (v7.310) with –auto (41). This alignment file was visualized using AliView (v1.28), and for receptor sequences, LRR regions were extracted to be inputted into Weblogo3 (v3.7.9) online server (42).

To assess CORE receptor variation, CORE gene sequences were extracted from available genomes from Benoit et al., using tblastN with default options and CORE from *S. lycopersicum* cv. Rio Grande as a reference (35). To build a phylogenetic tree, mafft was used to build a multiple sequence alignment (MSA) and iqtree2 (v2.1.2) was used for tree building using default options and 1000 ultrafast bootstraps (43). The Newark file was visualized using iTOL (44). Using the same MSA, normalized Shannon entropy was calculated in a positional context using a similar method as previously described (45). Detailed code for computational analyses can be found in the following GitHub repository (DanielleMStevens/mamp-prediction-ml).

### AlphaFold3 models

The AlphaFold3 online server was used to model the receptor ectodomain-protein ligand complex. The same receptor ectodomain sequence was used in both AlphaFold3 and mamp-ml. No additional modifications were made. Structural models were visualized using ChimeraX (v1.8). Code for visualization can be found in Zenodo (DOI: 10.5281/zenodo.15864665).

### Growth Conditions and measuring immunogenicity via ROS production

Solanaceous species characterized in this study can be found in Supplemental Table 6. These species were grown in growth chambers at 16/8-h light/dark cycles at 22.5°C for at least six to ten weeks after germination before measuring ROS burst. For measuring ROS, leaf punches were taken using cork borer no. 1 (diameter of 4 mm) from fully expanded leaves across biological replicates. Leaf punches (4 per plant) were floated on their abaxial side in 190 μl of sterile water in a 96-well white microtiter plate for 18 to 24 hours in the dark. To test ROS elicitation, sterile water was removed and a 100 μl solution of sterile water, L-012 (Wako Chemicals, Cat. No. 143556-24-5), horseradish peroxidase (HRP) (Sigma Aldridge, Cat. No. P6782), and 100 nM to 200 nM of peptide dissolved in sterile water or 100% DMSO. For flg22, 20 μM L-012 and 20 ug/mL HPR was used. For csp22, 40 μg/mL HPR was used. Assay measurements were taken every 1.5 minutes for a minimum 60 minutes. Luminescence was measured using an Infinite M200 Pro Luminescent Microplate Reader (Tecan Life Sciences, Switzerland). Across all ROS experiments, the maximum RLU was determined for each leaf punch and averaged between punches per plant. Peptides used in this study include those from Stevens et al. 2024 (7) and Li et al. 2025 (11) as well as those in Supplemental Table 7.

---

**Fig. S1.**
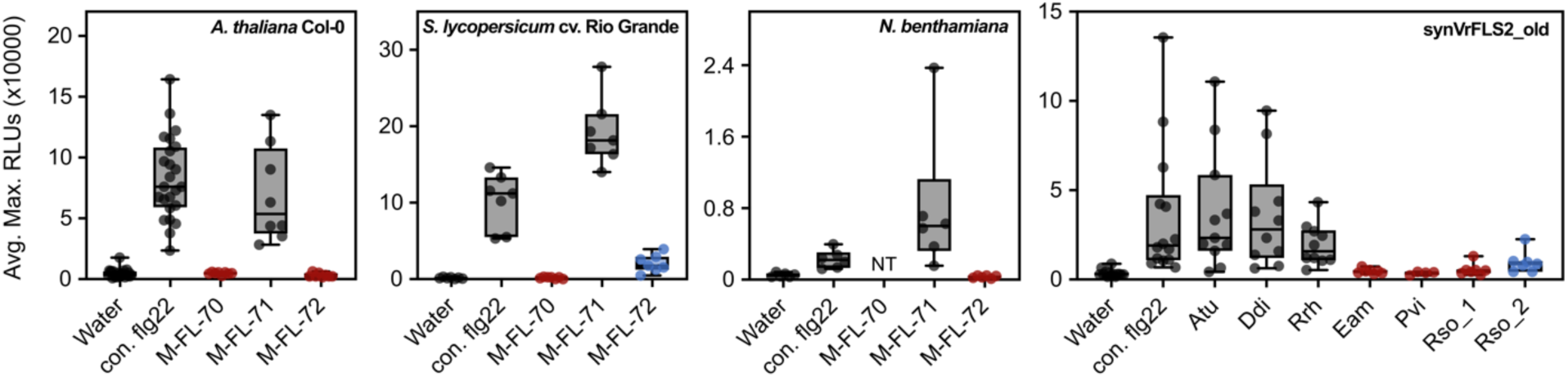
Additional training data collected across natural and synthetic FLS2-flg22 variants. Induction of ROS from flg22 variants across three species which encode a FLS2 homolog as well as one synthetic variant, synVrFLS2_old. 100 nM peptide was tested on at least three plants. Receptor-epitope interactions were categorized as the following: grey = immunogenic (around equal to or more than consensus flg22), red = non-immunogenic (equal to water), and blue = weakly immunogenic (induction of ROS but notably less than consensus flg22. NT = Not tested.

**Fig. S2.**
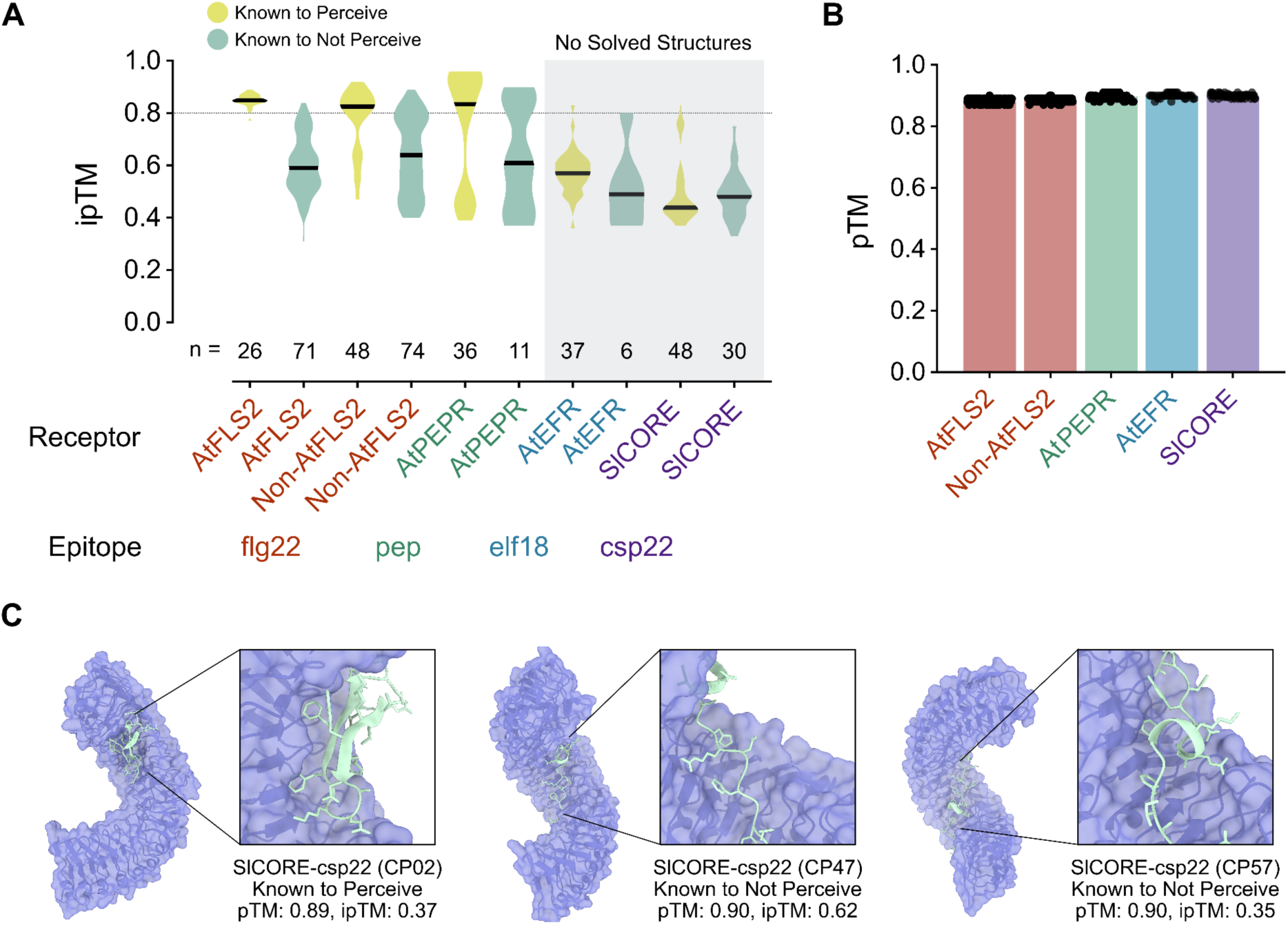
AlphaFold3 fails to predict recognition using ipTM scores when an underlying solved structure is not present. **(A-B)** Using AlphaFold3 metrics, ipTM (left) and pTM (right), for perception based on those that were experimentally determined across several published studies. FLS2-flg22 data is from Li et al., 2024 (11), EFR-elf18 data is from Sutherland et al., 2025 (3), and CORE-csp22 and PEPR-pep is from this work. At = Arabidopsis thaliana Col-0, Sl = Solanum lycopersicum (tomato) cv. Rio Grande. **(C)** Examples of AlphaFold3 models from CORE-csp22 variants that have been experimentally verified. Structures were visualized using ChimeraX.

**Fig. S3.**
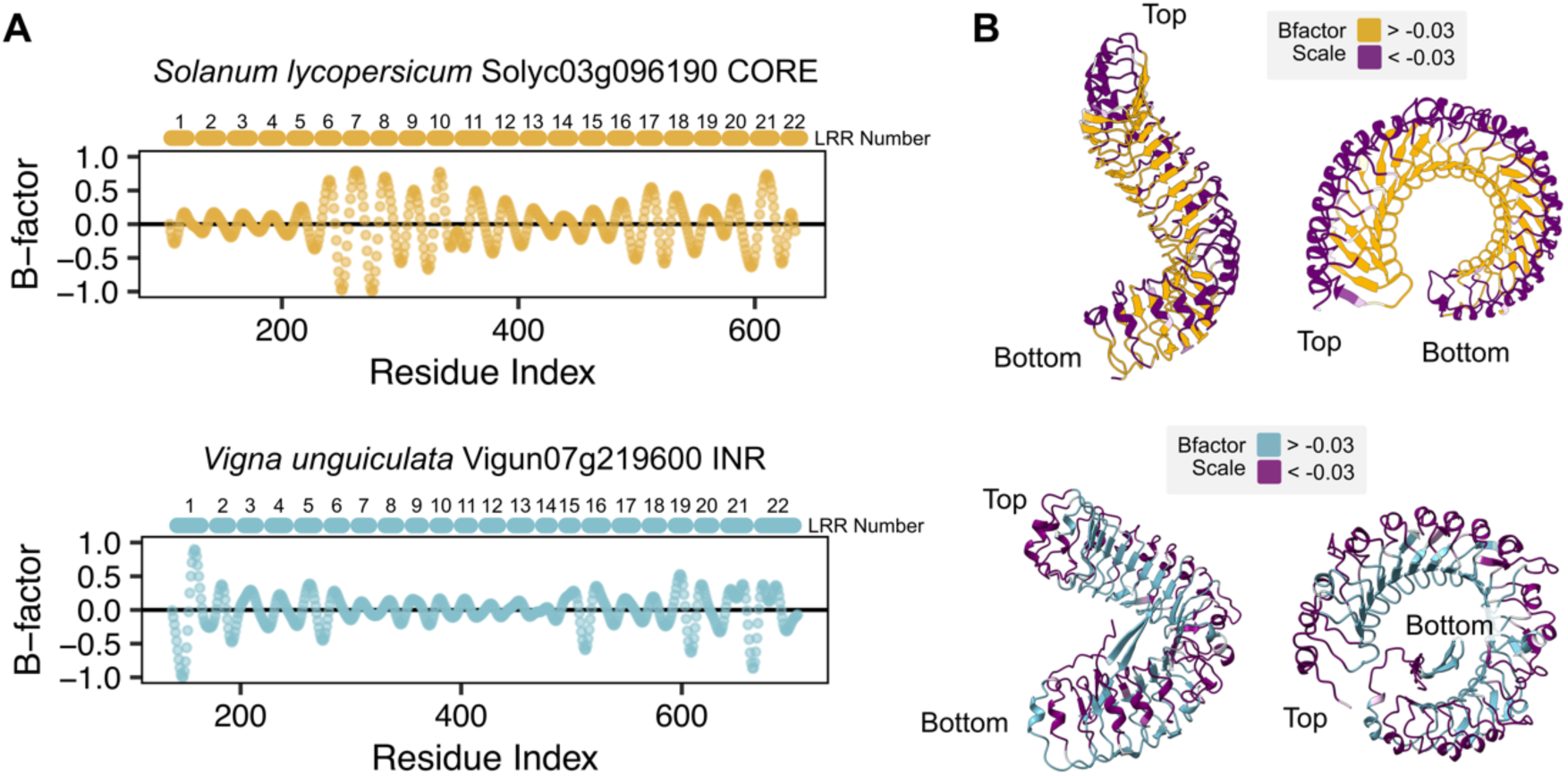
Using bandpass B-factor values from LRR-Annotation as a metric to count LRR repeats and capture the inner concave surface. **(A)** Bandpass b-factor values are extracted which represent each LRR as a repeating interval. Two examples where LRR-Annotation predicts the correct (top) and incorrect (bottom) number of LRRs based on bandpass B-factor transitions across the x-axis. **(B)** Bandpass b-factor values plotted on associated ectodomain structures where positive b-factor values represent the inner concave surface and negative b-factor values represent the outer receptor surface. Structures were generated using AlphaFold3 and visualized using ChimeraX.

**Fig. S4.**
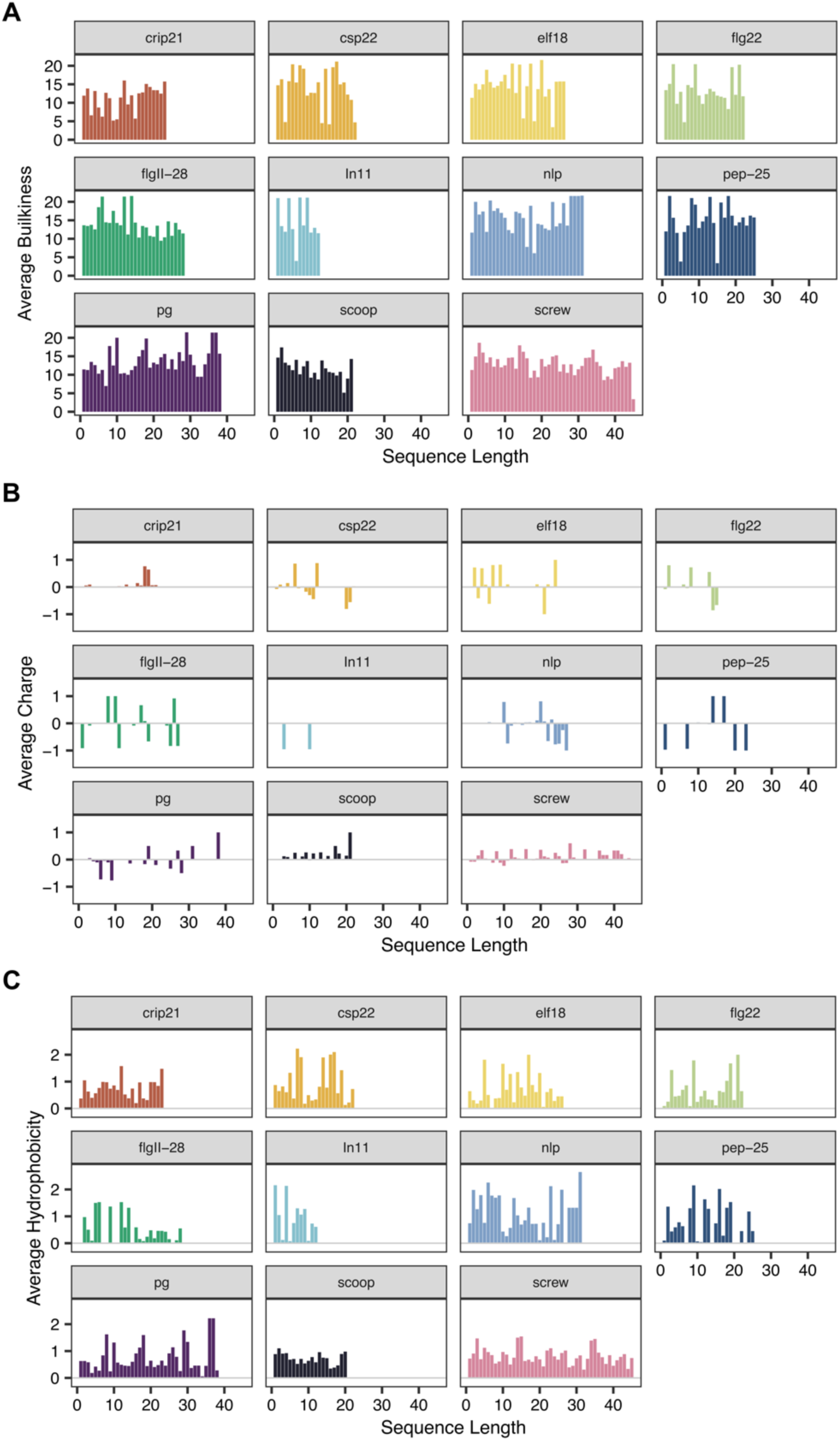
Amino acid side chain properties generate distinguishable features between epitopes. Average bulkiness **(A)**, charge **(B)**, and hydrophobicity by the Manavalan scale **(C)** for each protein ligand in a positional context.

**Fig. S5.**
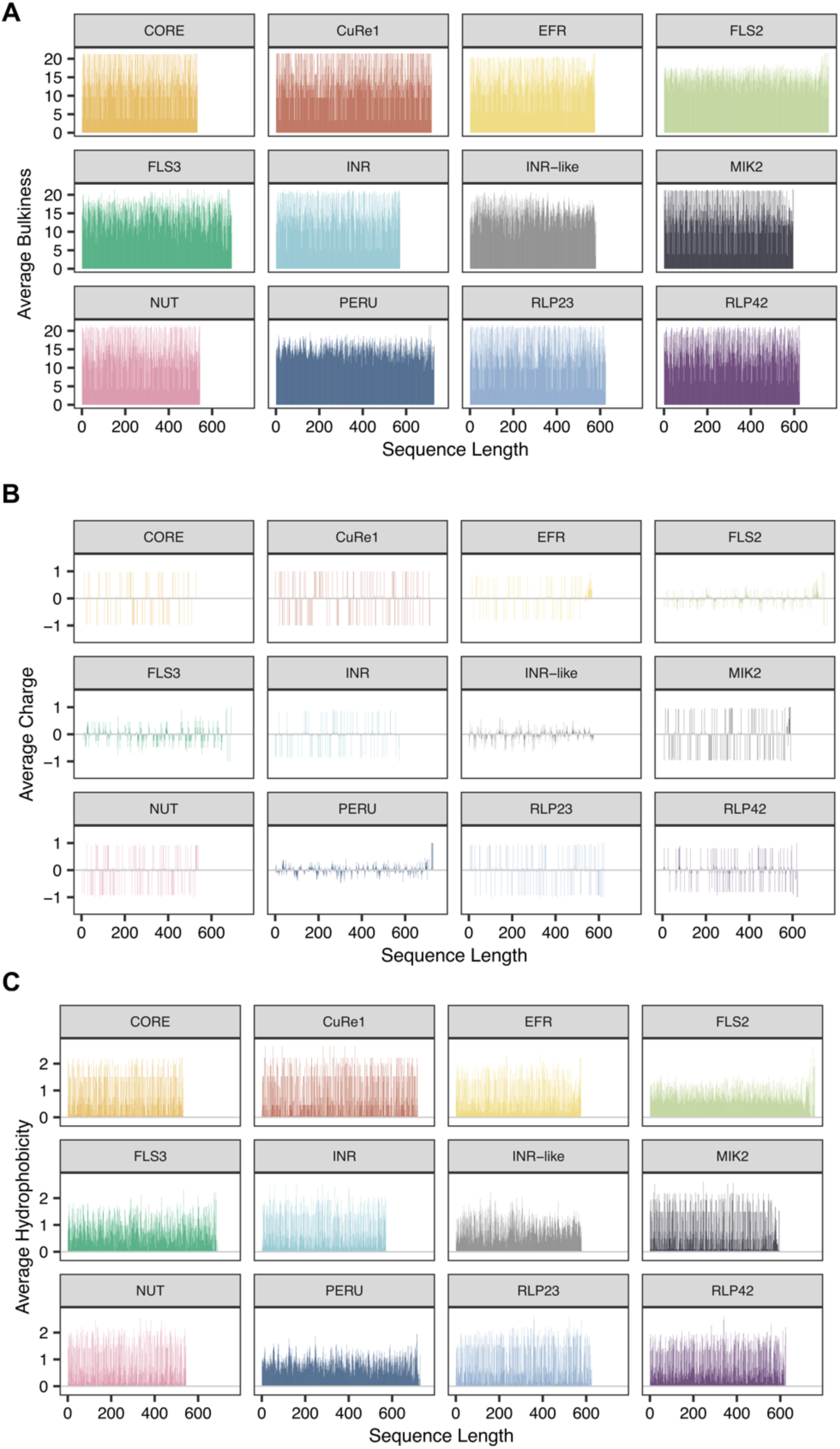
Amino acid side chain properties enable distinguishable features between receptor ectodomains. Average bulkiness **(A)**, charge **(B)**, and hydrophobicity by the Manavalan scale **(C)** for each protein receptor ectodomain in a positional context.

**Fig. S6.**
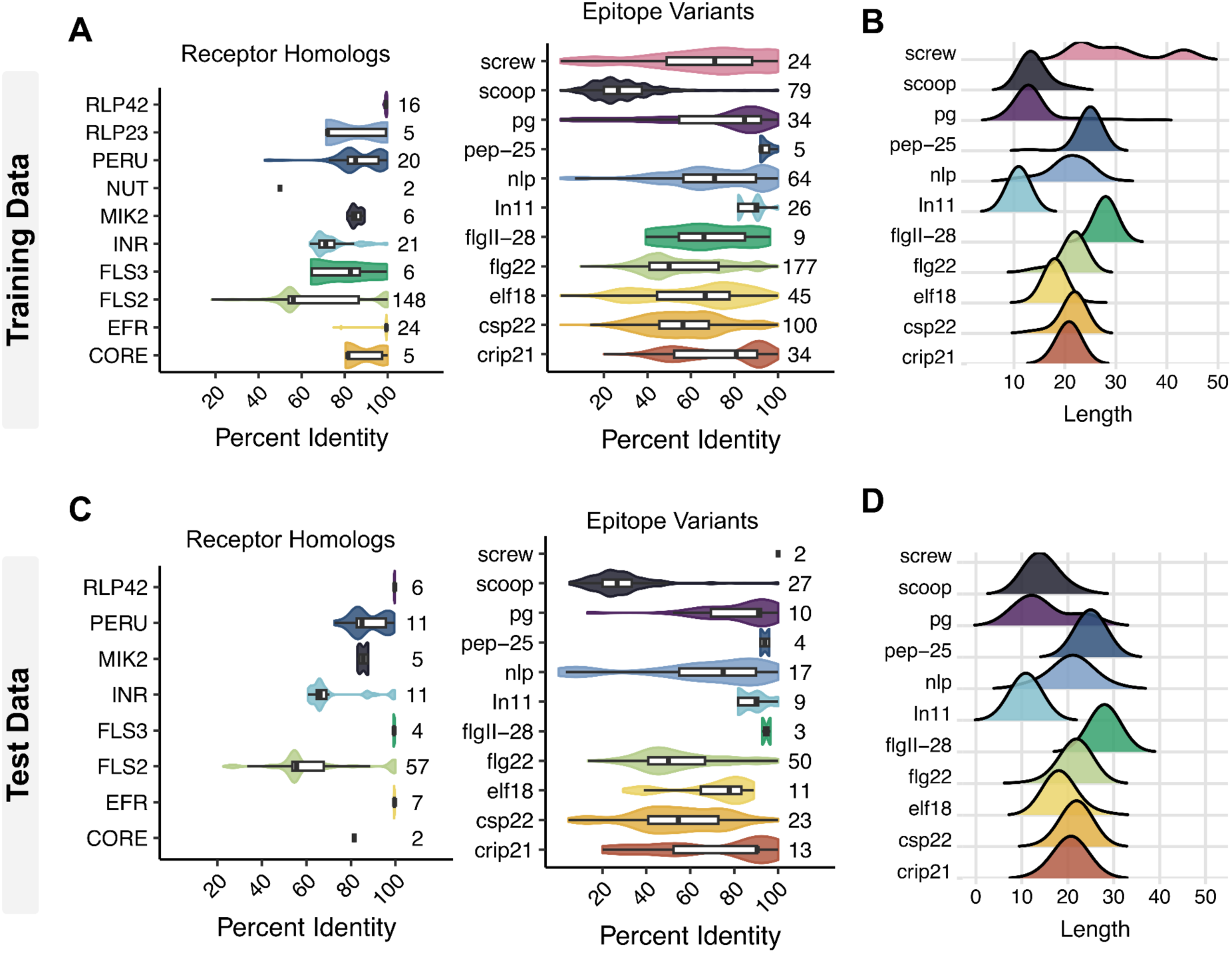
Sequence diversity of training and test data for both receptors and their protein ligands. Sequence diversity **(A)** and ligand length **(B)** for training data. Sequence diversity **(C)** and ligand length **(D)** for test data.

**Fig. S7.**
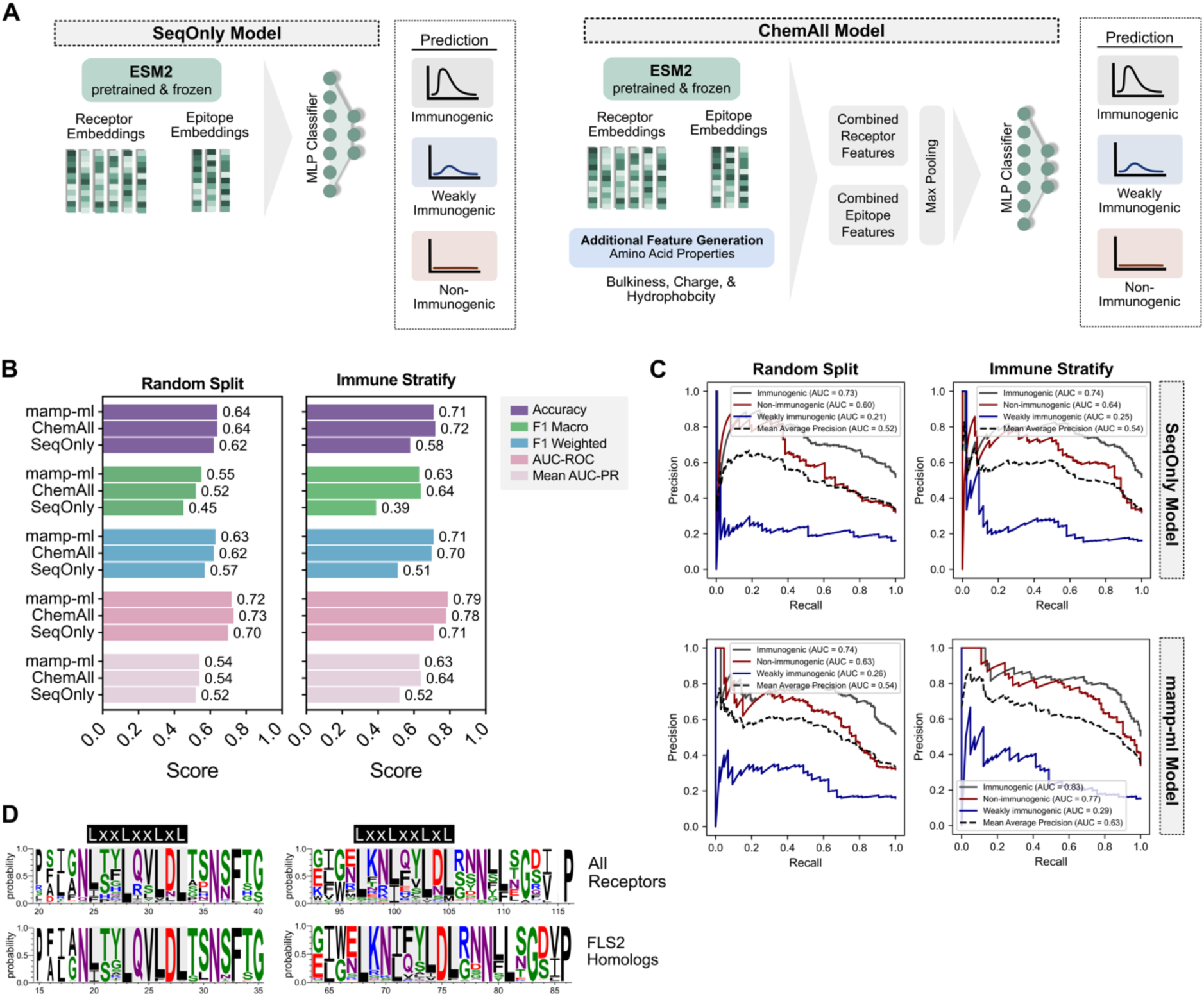
Performance metrics under different training data preparations and model architectures. **(A)** Diagram of current model architecture of mamp-ml versus two baseline models, one of which relies only on sequence embeddings (SeqOnly) and one which combines sequence embeddings with chemical features (ChemAll) **(B)** Model accuracy and performance of mamp-ml, SeqOnly, and ChemAll using a stratified split by immunogenicity (right) and a completely randomized split, which shuffles the receptor and epitope sequences for the training data (left). For all three models, the same hyperparameters including learning rate (0.0004), batch size (8), and epoch number (20) were used. **(C)** AUPRC of validation data for SeqOnly and mamp-ml when stratifying by immunogenicity (right) and a completely randomized split (left). **(D)** Weblogos along LRR domain sequence from all receptors and FLS2 homologs.

**Fig. S8.**
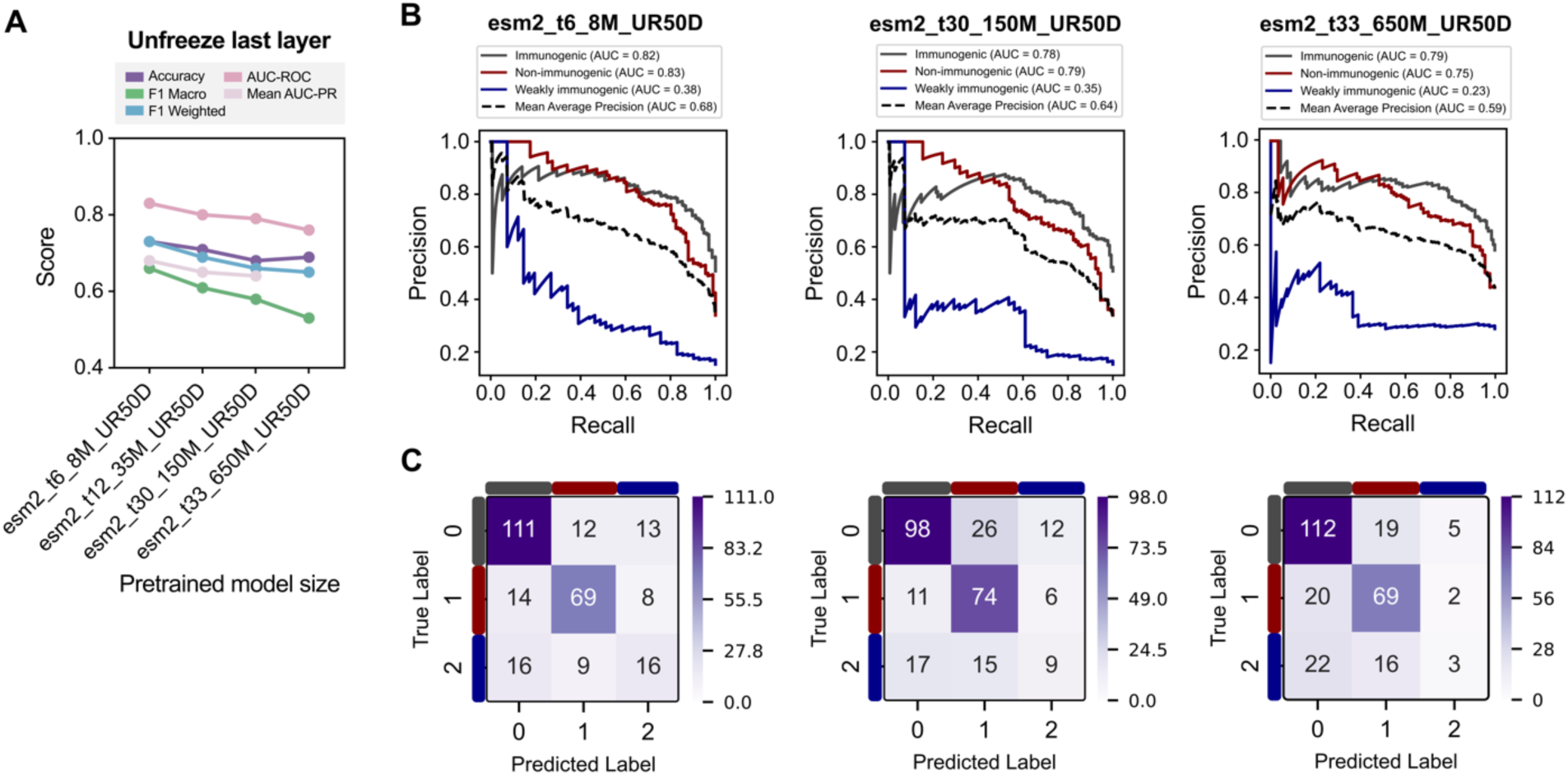
Larger ESM-2 pretrained models led to overfitting and reduced prediction accuracy. For all four model versions, the same hyperparameters including learning rate (0.0004), batch size (8), and epoch number (20) were used. **(A)** Model accuracy and performance of mamp-ml using different-sized ESM-2 pretrained models. **(B)** AUPRC of test data across three model sizes. **(C)** Confusion matrix of validation data across three model sizes. Labels are the following: 0 = immunogenic, 1 = non-immunogenic, and 2 = weakly immunogenic.

**Fig. S9.**
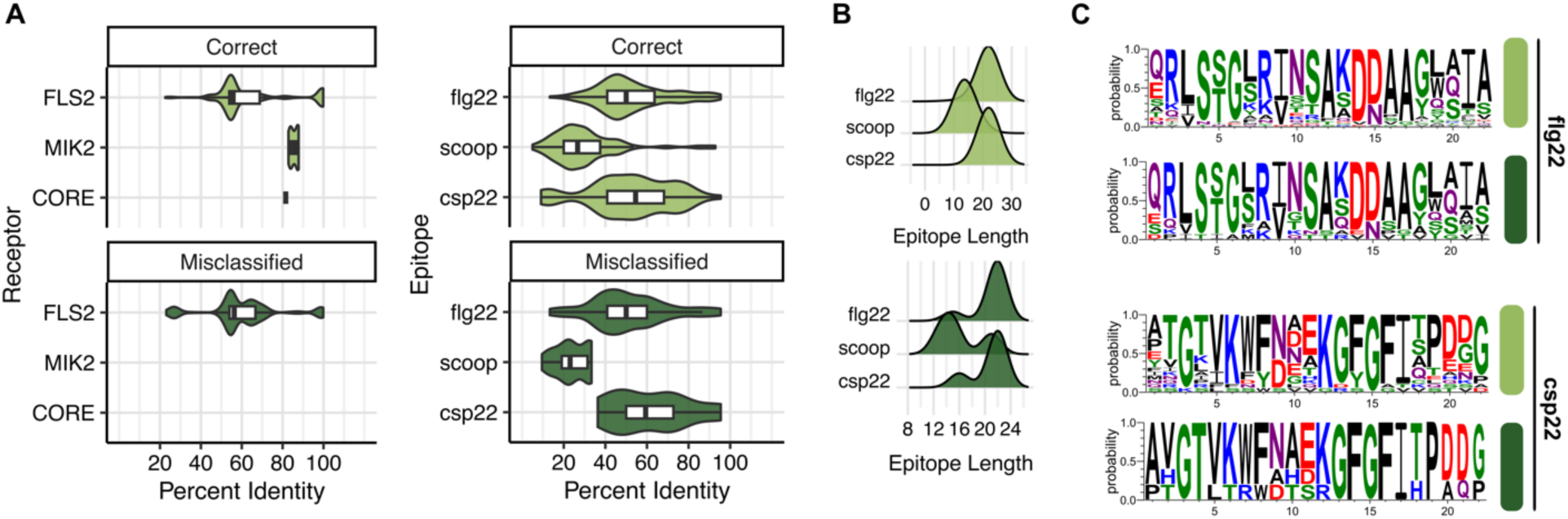
Example of receptor-epitope combinations that are misclassified in the test dataset. **(A)** Global all-by-all sequence similarity of receptors (left) and epitopes (right) of those that were correctly predicted and misclassified. **(B)** Ligand length of epitopes that were correctly predicted and misclassified. **(C)** Weblogos of epitopes that were correctly predicted and misclassified (flg22: top, csp22: bottom). Light green and dark green colors signify correct and misclassified predictions, respectively.

**Fig. S10.**
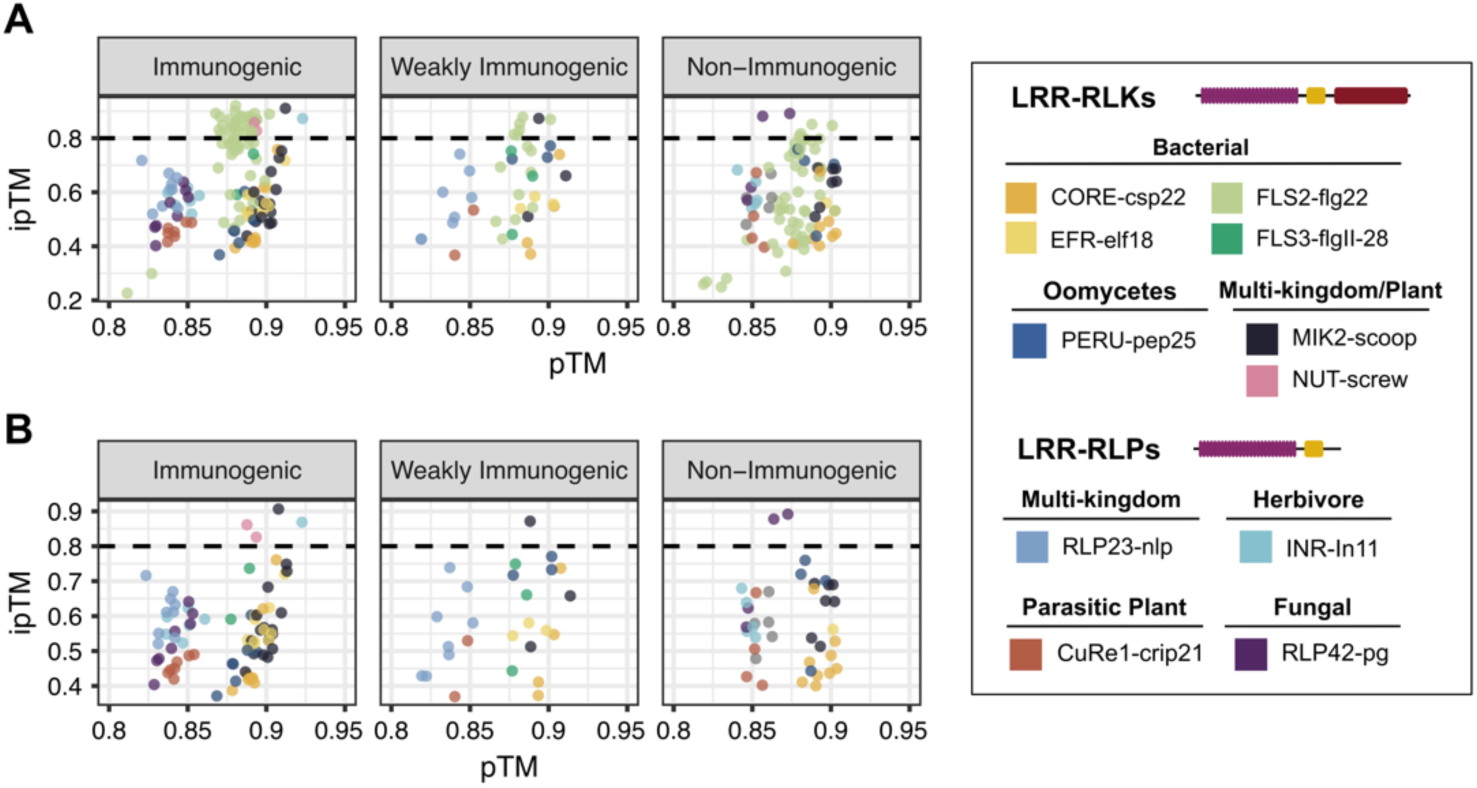
AlphaFold3 metrics on test data. Global prediction accuracy (predicted template modeling: pTM) and protein-protein interactions (interface predicted template modeling: ipTM) of the test dataset with **(A)** and without **(B)** FLS2-flg22 combinations. The dashed line at 0.8 ipTM represents those protein-protein interaction scores with high confidence.

**Fig. S11.**
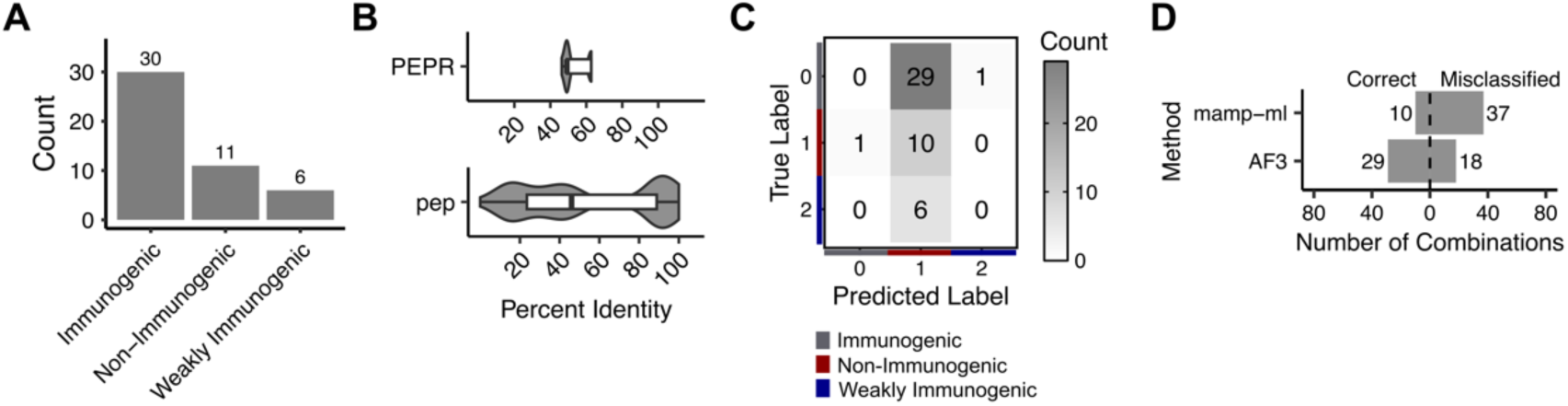
Assessing model accuracy using unseen receptor-epitope interaction, PEPR-pep. **(A)** Distribution of receptor-epitope interaction data (n = 47) based on their immunogenic outcomes collected for model assessment. **(B)** Receptor ectodomain (top) and epitope (bottom) sequence diversity based on all-by-all percent identity comparisons. **(C)** Confusion matrix of mamp-ml prediction. Labels are the following: 0 = immunogenic, 1 = non-immunogenic, and 2 = weakly immunogenic. **(D)** Summary of correct and misclassified predictions between mamp-ml and AlphaFold3 using an ipTM score cutoff of 0.8.

**Fig. S12.**
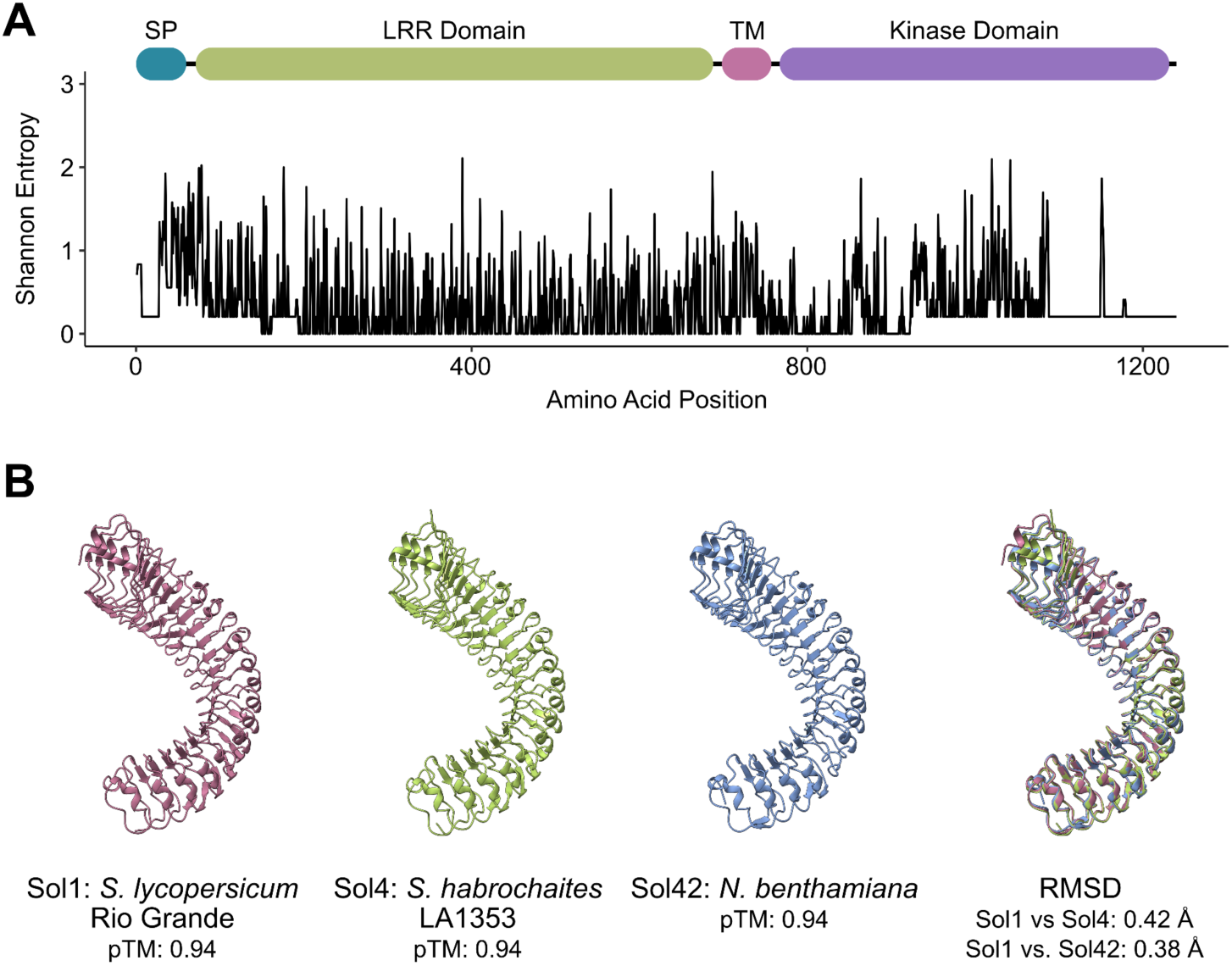
Structurally conserved but sequentially diverse CORE receptor variants. **(A).** Shannon entropy of 16 CORE homologs in respect to protein domain structure. **(B)** Structural models of CORE receptors ectodomain used model testing in Fig. 4C-D. Models were predicted via AlphaFold3 and visualized using ChimeraX.

**Fig. S13.**
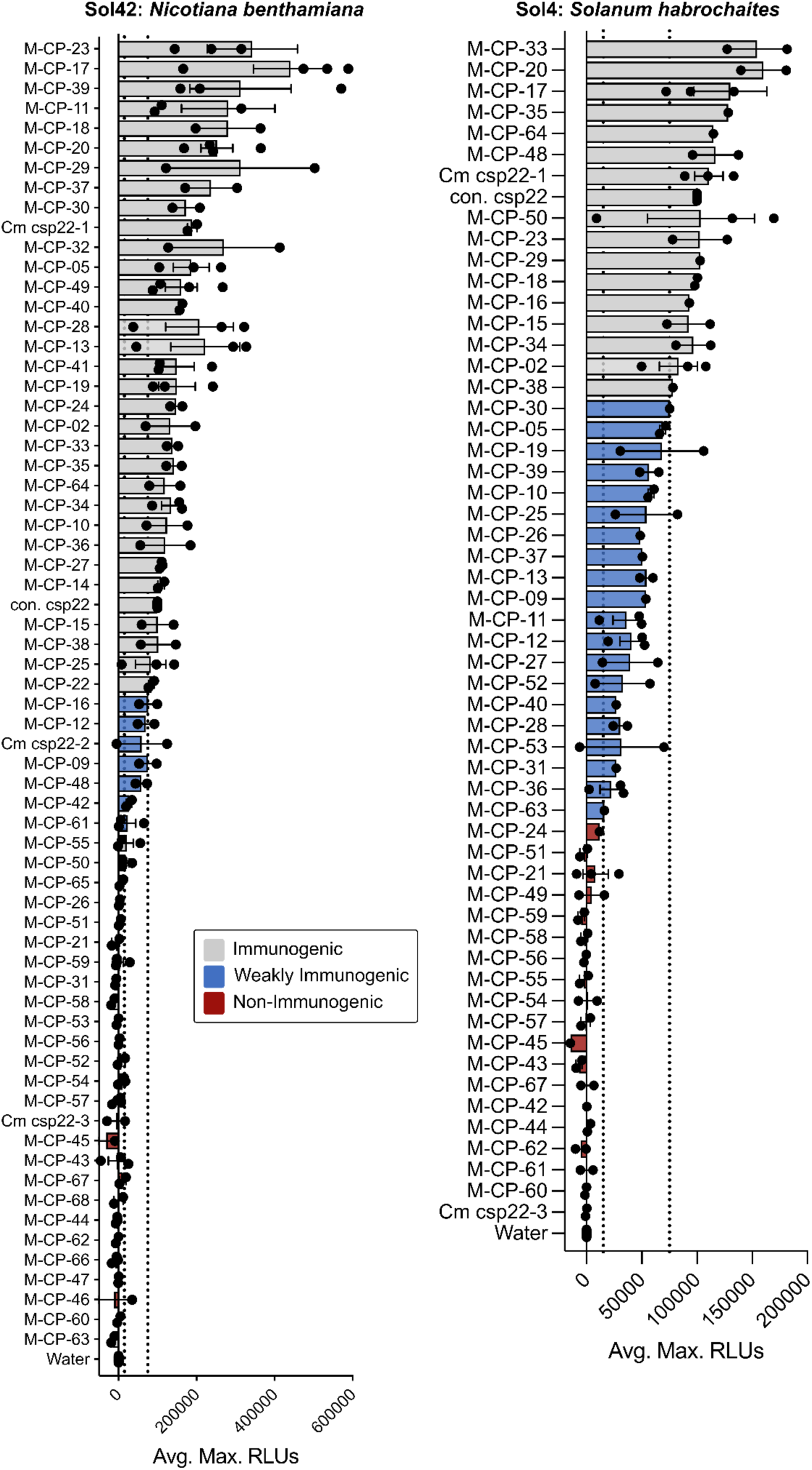
Variable CORE-csp22 sequences influence immunogenic outcomes. Categorization of immunogenic outcomes based on ROS magnitude (immunogenic: ∼75,000+ RLUs, weakly immunogenic: ∼15-75,000 RLUs, and non-immunogenic: >15,000 RLUs). 200 nM peptide was tested with at least three plants per replicate.

**Fig. S14.**
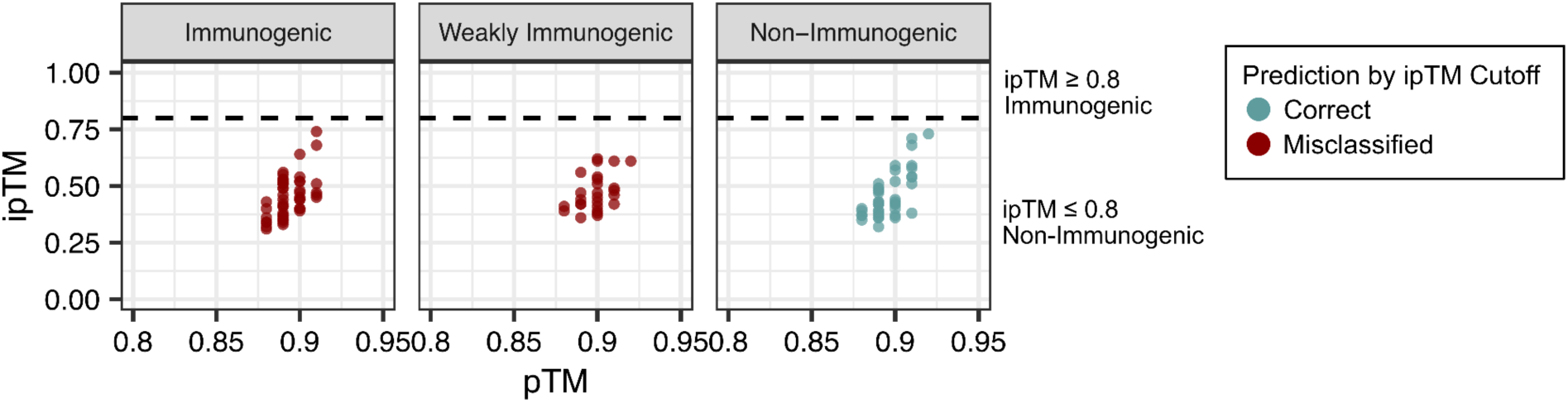
AlphaFold3 metrics on validation data. Global prediction accuracy (predicted template modeling: pTM) and protein-protein interactions (interface predicted template modeling: ipTM) of the validation dataset. The dashed line at 0.8 ipTM represents those protein-protein interaction scores with high confidence.

**Fig. S15.**
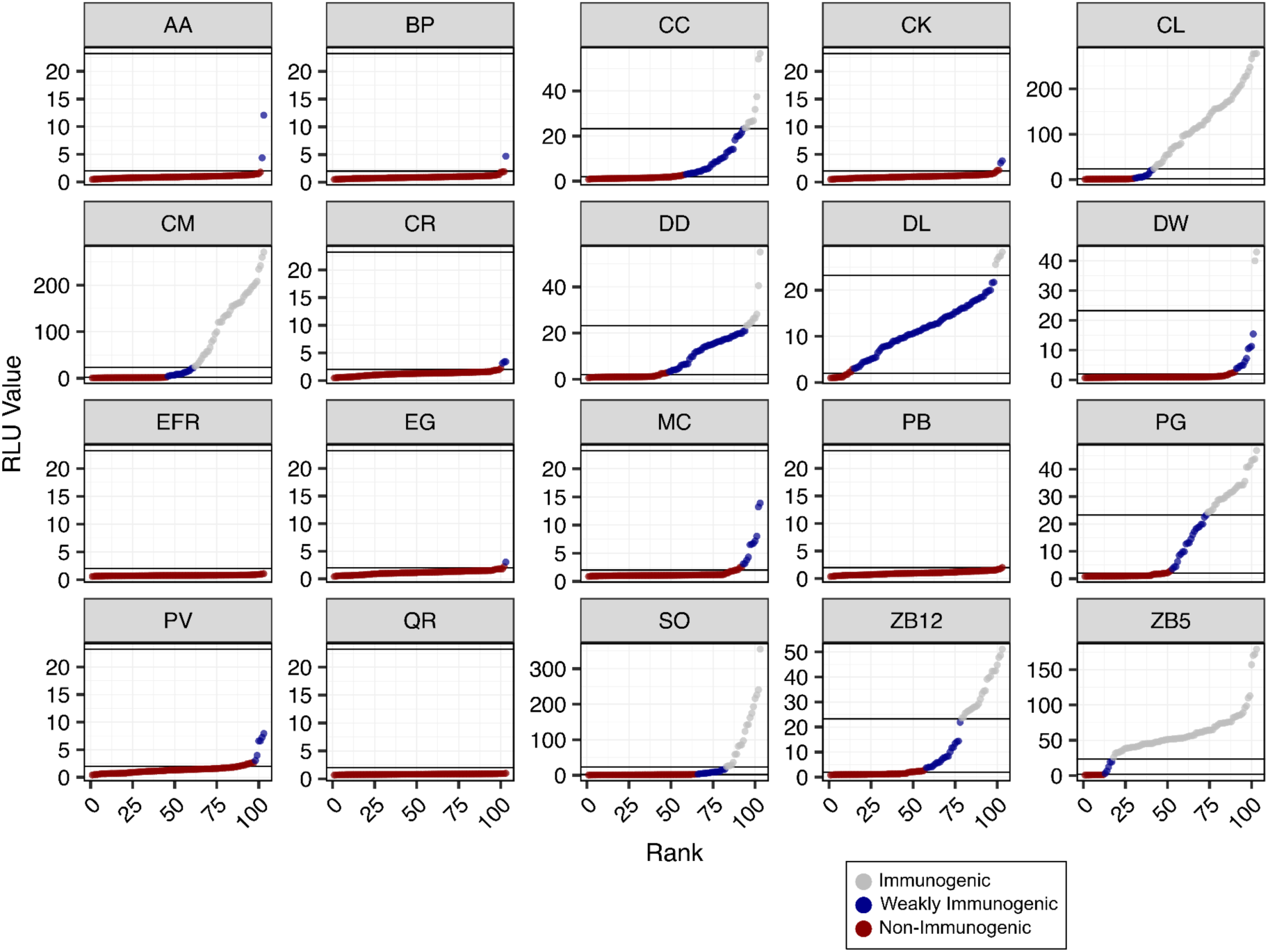
SCORE orthologs from Ngou et al. categorized by immunogenicity. Reactive Oxygen Species (ROS) production of each receptor ortholog and csp15 ligand variant ranked by their Reactive Light Unit (RLU) values. Those RLUs below 3, as denoted in Ngou et al. 2025, are categorized as non-immunogenic. Those RLUs above 3 are classified as immunogenic or weakly immunogenic based on the relative maximum RLU value.

**Fig. S16.**
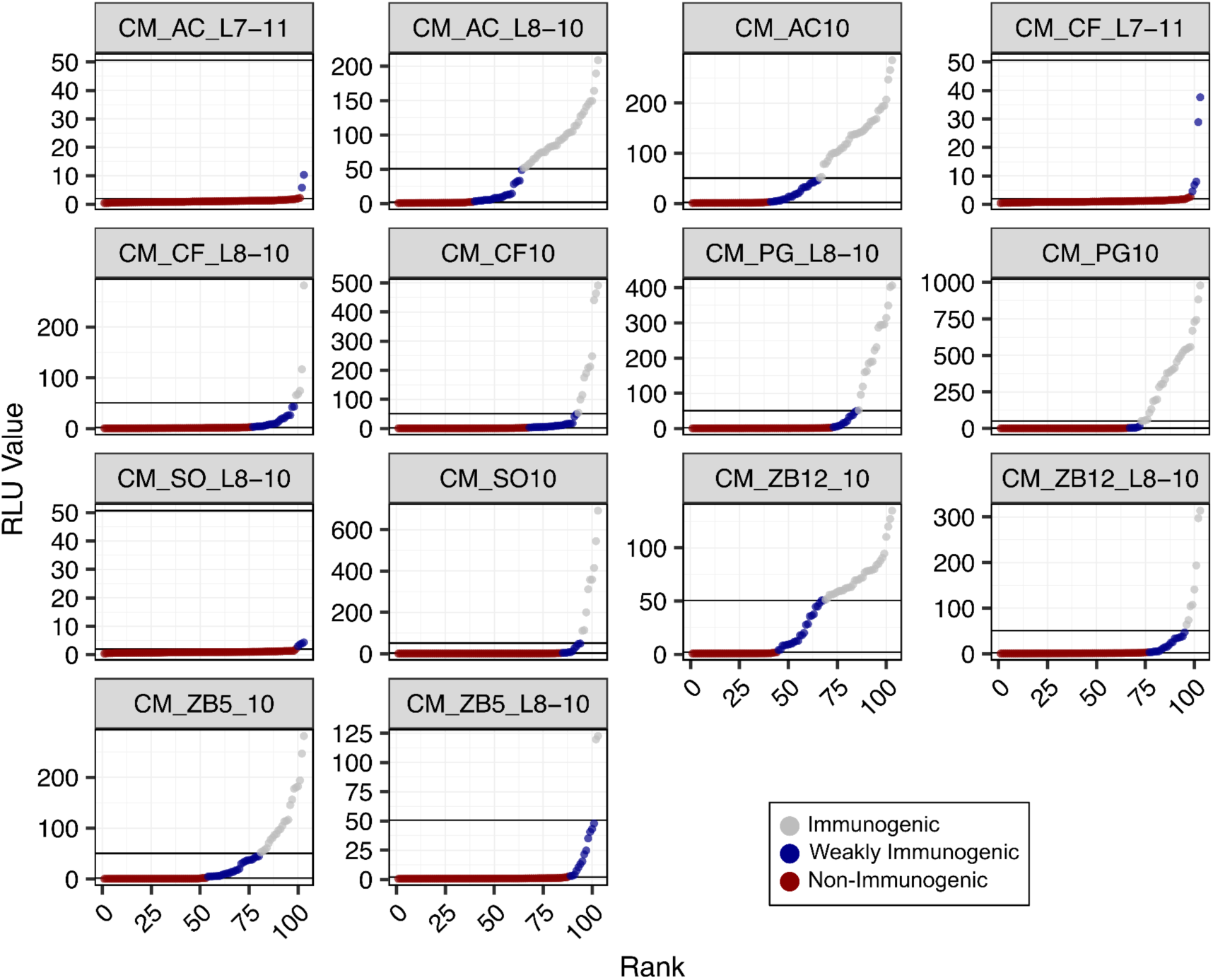
SCORE chimeras from Ngou et al. categorized by immunogenicity. Reactive Oxygen Species (ROS) production of each receptor chimera and csp15 ligand variant ranked by their Reactive Light Unit (RLU) values. Those RLUs below 3, as denoted in Ngou et al. 2025, are categorized as non-immunogenic. Those RLUs above 3 are classified as immunogenic or weakly immunogenic based on the relative maximum RLU value.

**Fig. S17.**
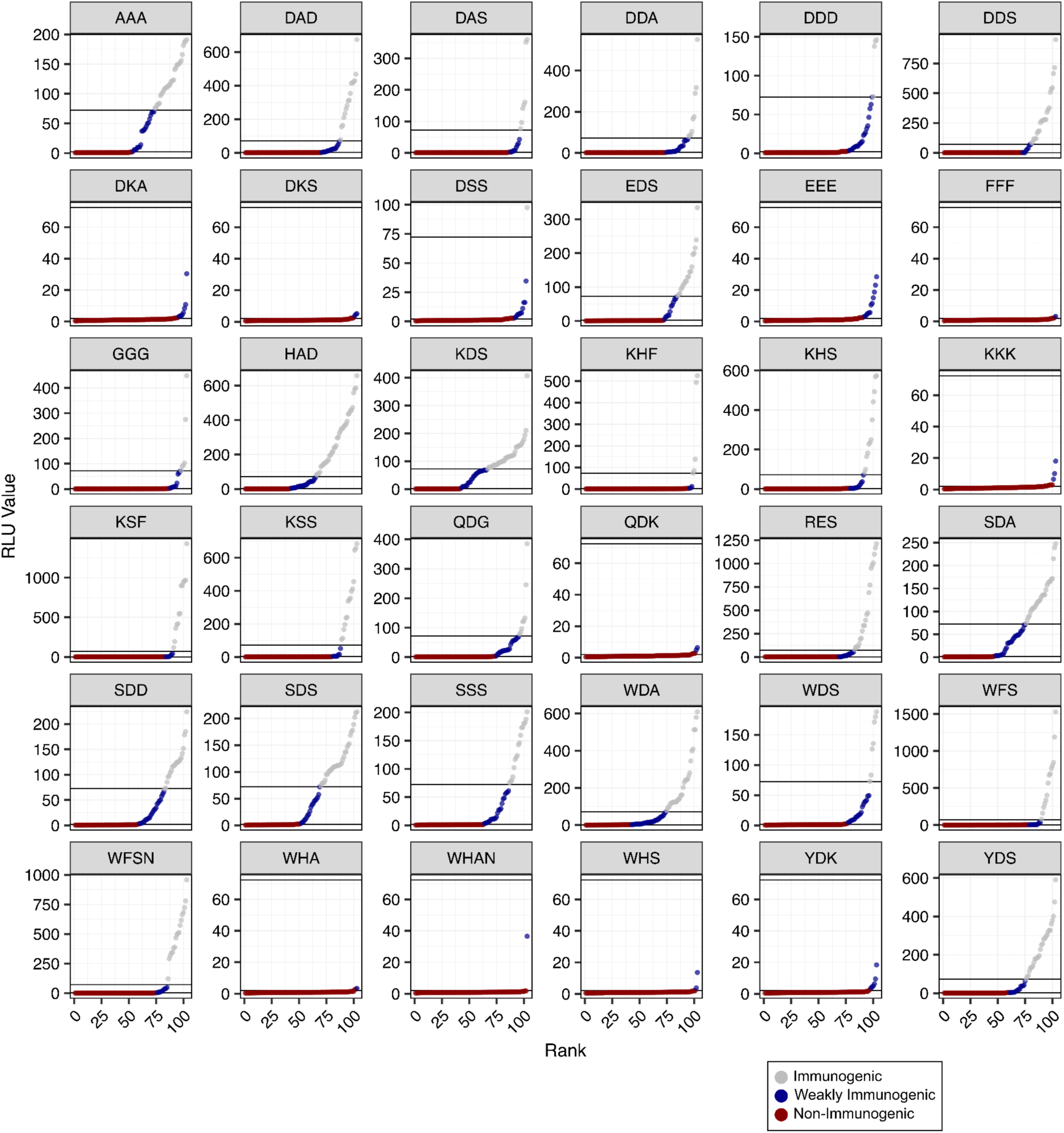
Amino acid substitution SCORE variants from Ngou et al. categorized by immunogenicity. Reactive Oxygen Species (ROS) production of each receptor variant and csp15 ligand variant ranked by their Reactive Light Unit (RLU) values. Those RLUs below 3, as denoted in Ngou et al. 2025, are categorized as non-immunogenic. Those RLUs above 3 are classified as immunogenic or weakly immunogenic based on the relative maximum RLU value.

**Fig. S18.**
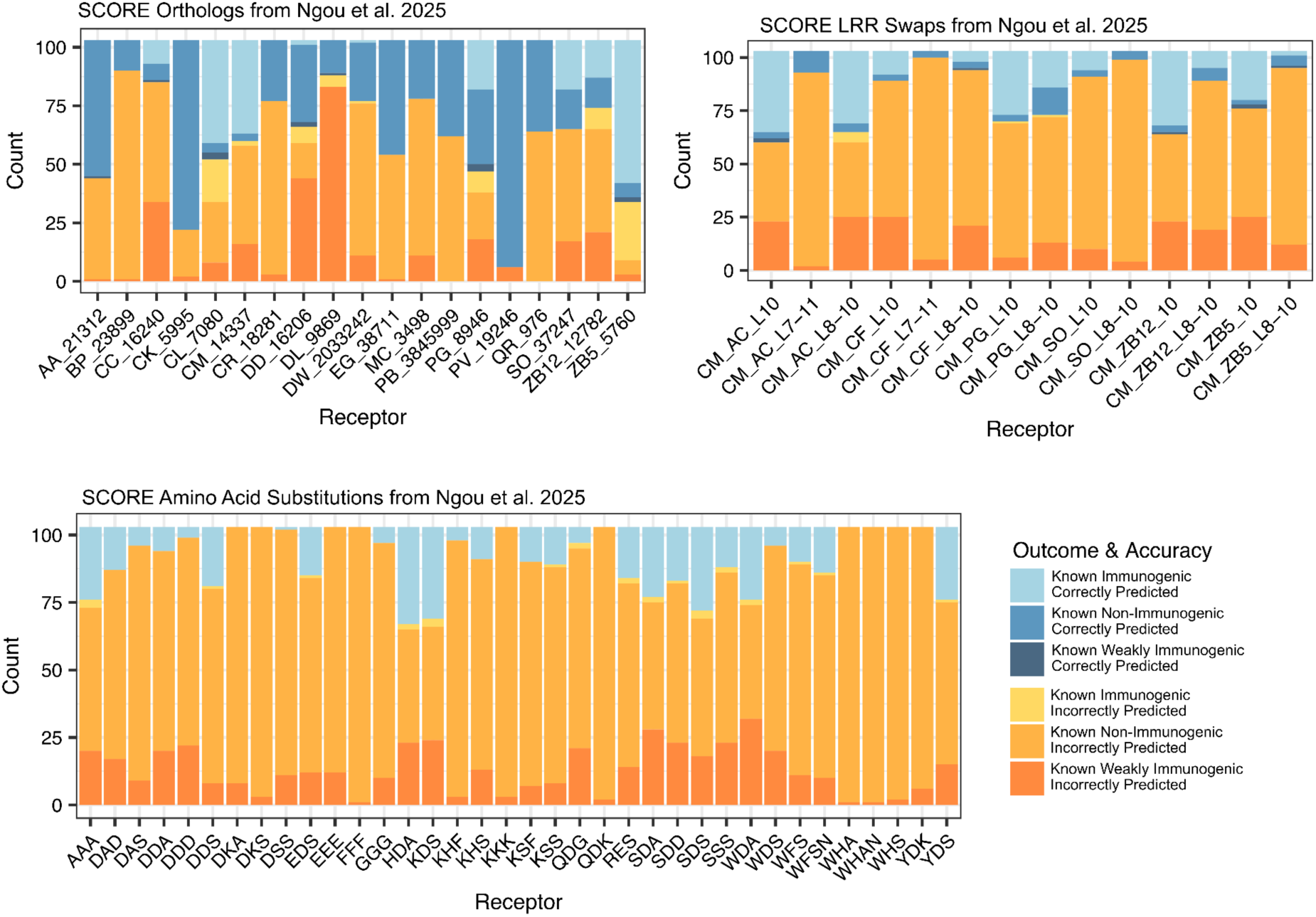
Zero-shot immunogenicity prediction of SCORE-csp22 variants. SCORE variants from Ngou et al. 2025 (13) were categorized into three groups: orthologs, chimeras (LRR swaps), and single point mutations (amino acid substitutions). Using mamp-ml, we zero-shot predicted SCORE-csp22 immunogenic outcomes across 103 csp22 variants and categorized by their known outcomes and prediction accuracy.

